# *In situ* Proteomics Unveils Specialized Domains for Extrasynaptic Signaling on Neuronal Cilia

**DOI:** 10.1101/2025.09.10.675452

**Authors:** Chia-Hsiang Chang, Van Ngu Trinh, Nidhi Rani Lokesh, Catalina Kretschmar Montecinos, Mark E. Pownall, Marian Kalocsay, Maxence V. Nachury

**Affiliations:** Department of Ophthalmology, University of California, San Francisco, CA 94143, USA; Smith Cardiovascular Research Institute, University of California San Francisco; CA 94143, USA; Department of Experimental Radiation Oncology, The University of Texas MD Anderson Cancer Center, Houston, TX 77030, USA; Department of Biochemistry and Biophysics, University of California, San Francisco, San Francisco, CA 94143, USA

## Abstract

Neuronal cilia have emerged as crucial signaling hubs, yet their molecular composition and integration with synaptic communication remain poorly understood. Using a newly developed *Arl13b-TurboID* mouse model, we achieved robust cilia-specific biotinylation and proteomic profiling across diverse tissues and cell types. Comparative proteomics revealed striking tissue-specific specialization, with neuronal cilia uniquely enriched in synaptic proteins, adhesion molecules, and neurotransmitter receptors. Surprisingly, several signaling and adhesion molecules localize to neuronal cilia in discrete nanodomains maintained by active retrieval mechanisms. In the mouse cortex, expansion microscopy revealed that the NMDA receptor subunit GluN1 is organized in nanodomains on neuronal ciliary membranes, which are precisely positioned to sample neurotransmitter efflux from neighboring glutamatergic synapses. These findings establish neuronal cilia as specialized extrasynaptic signaling platforms, with nanoscale organization enabling them to integrate local synaptic cues and modulate neuronal connectivity.

## INTRODUCTION

Primary cilia are micron-scale cellular projections that receive and transduce signals^1,2^. In the central nervous system (CNS), most neurons and glial cells harbor a primary cilium, and human genetic studies have linked ciliary dysfunction to neurodevelopmental and cognitive disorders^3^, including Bardet-Biedl syndrome (BBS)^4^, Joubert syndrome^5^, schizophrenia^3,6–8^, bipolar disorder^3,6^, autism spectrum disorders^9–13^ and Parkinson’s disease^14–17^. Despite their emerging role in neural signaling, the molecular mechanisms by which neuronal cilia contribute to brain connectivity and communication remain poorly understood.

Neuronal cilia are increasingly recognized as extrasynaptic devices that mediate communication between neurons and other cells within the brain^18,19^, acting as cellular antennae that transduce signals^20,21^. In specialized sensory neurons, such as photoreceptors and olfactory receptor neurons, cilia organize entire signaling cascades^22,23^. By contrast, the function of cilia in most brain cells remains poorly understood. Recent studies suggest that neuronal cilia may influence synaptic connectivity^24–26^. Volume electron microscopy has revealed that neuronal cilia are integrated into the neuronal circuitry in the mammalian brain^27–29^: mouse hippocampal cilia form intimate associations with axons named axociliary synapses^27^, human cortical cilia frequently contact tripartite synapses involving astrocytes^29^, and mouse visual cortex cilia are positioned to eavesdrop on neighboring synaptic signals^28^. Despite these advances, the molecular underpinnings of cilia-synapse interactions remain elusive.

Proteomic studies of sensory cilia have been facilitated by their biochemical isolation, enabling detailed characterization of their molecular makeup^30–33^. However, most neuronal cilia cannot be purified, leaving their proteome largely unexplored. To address this gap, we developed an *in situ* proximity labeling approach in mice and systematically profiled the brain cilia proteome. We discovered that neuronal cilia harbor a diverse array of signaling proteins, adhesion molecules, and neurotransmitter receptors, distinct from renal cilia. Using expansion microscopy, we reveal the nanoscale spatial organization of ciliary proteins and demonstrate that neuronal cilia are closely positioned to glutamatergic synapses. Notably, the NMDA receptor subunit GluN1 is non-randomly distributed on ciliary membranes near synaptic structures, suggesting that cilia may ‘eavesdrop’ on synaptic communication.

## RESULTS

### ARL13B-TurboID enables efficient and cilium-selective biotinylation in mice

To determine the most efficient proximity labeling strategy in neuronal cilia, we screened various ciliary targeting signals and proximity labeling enzymes via lentiviral transduction of cultured cerebellar granule neuron progenitors (GNPs) (**Fig. S1A**). While well-validated in other systems, the ciliary targeting sequence of NPHP3^34^, and the ciliary GPCR somatostatin receptor 3 (SSTR3) were only partially enriched in GNP cilia (**Fig. S1B-C**). In contrast, ARL13B efficiently and specifically targeted GNP cilia (**Fig. S1B-C**). Surprisingly, despite achieving robust biotinylation of the cilia proteome in kidney cells^34,35^, the ascorbate peroxidase APEX2 fused to ARL13B failed to biotinylate the ciliary content of GNPs, even under elevated biotin-phenol concentrations (**Fig. S1D-E**). Instead, the engineered biotin ligase TurboID^36^ carried out efficient and selective biotinylation in GNPs cilia when fused to ARL13B (**Fig. S1D-E**). Similarly, ARL13B-TurboID efficiently biotinylated ciliary contents in multiciliated ependymal cells (**Fig. S1F-G**), and in the human cerebellar Daoy cell line (**Fig. S1H-I**). The V359A mutation of the ciliary targeting motif of human ARL13B^37^ abolished ciliary biotinylation, confirming the specificity of the system (**Fig. S1H-I**). Collectively, these findings demonstrate that ARL13B-TurboID achieves robust, cilium-specific biotinylation across various CNS cell types.

To implement cilia-specific biotinylation *in vivo* while avoiding overexpression-related perturbations^38–40^, we inserted a *GFP-TurboID* cassette at the endogenous *Arl13b* locus to express ARL13B-GFP-TurboID (referred to as ARL13B-TurboID) (**Fig. 1A**). The *Arl13b-GFP-TurboID* allele did not affect development, body weight or cilia length in heterozygous and homozygous state (**see Fig. S2F, S5A-B**), indicating that the fusion protein is fully functional. We next introduced a point mutation in the ciliary targeting signal of mouse ARL13B^37^ to generate the *Arl13b[V358A]-TurboID* allele (**Fig. 1A**).

**Figure 1.**
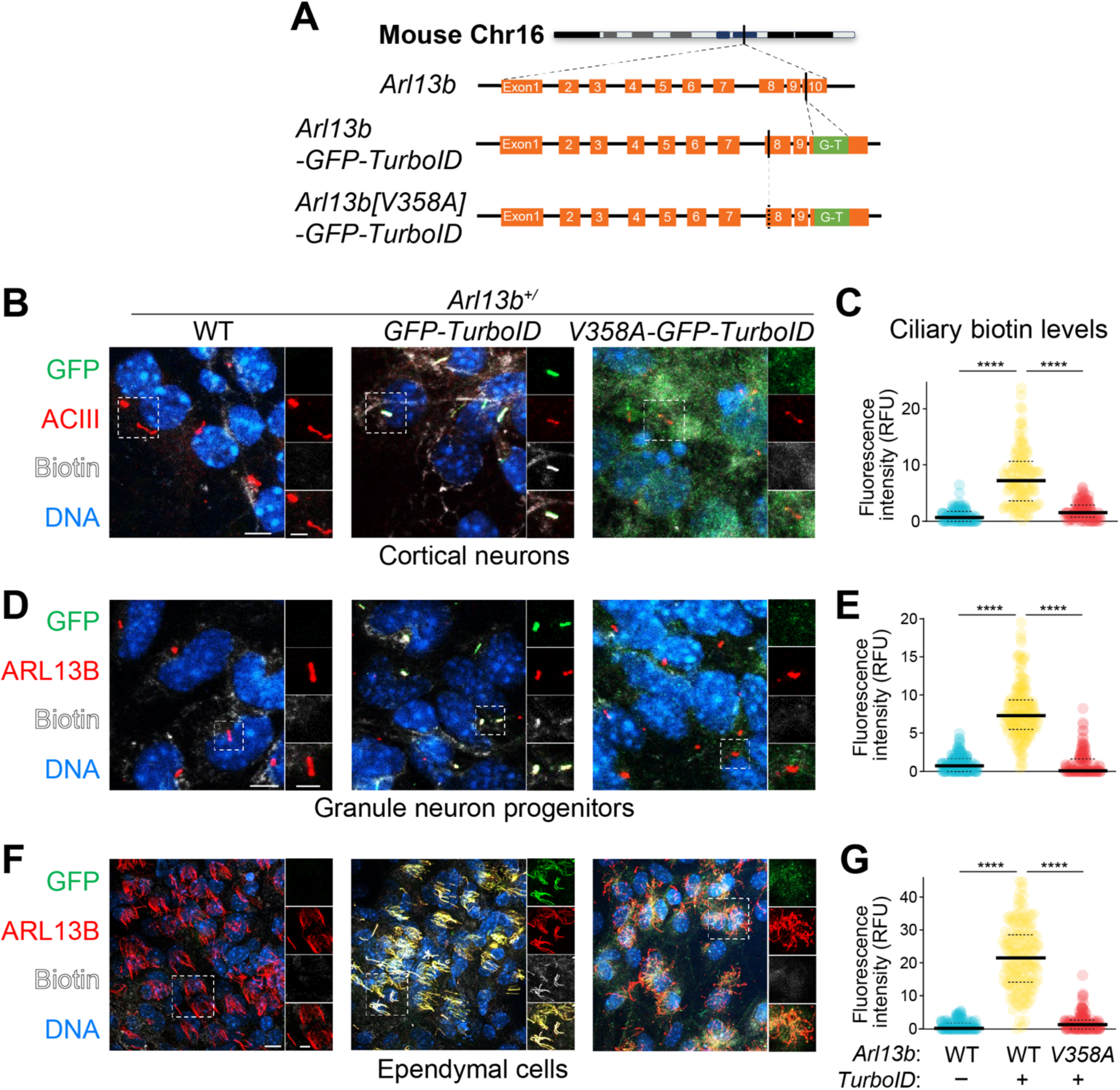
ARL13B-TurboID robustly and specifically biotinylates ciliary proteins. **A.** Diagram of the targeting strategy. The GFP-TurboID (G-T) cassette was inserted into exon 10 of *Arl13b* in the mouse genome immediately before the stop codon to generate the *Arl13b-GFP-TurboID* allele. Mutagenesis at Exon 8 generated the *Arl13b[V358A]-GFP-TurboID* allele. **B-G.** Cilium-specific biotinylation by ARL13B-TurboID. Cortical neurons **(B)**, granule neuron progenitors (GNPs, **D**), and ependymal cells **(F)** were cultured from E15.5 cerebral cortex, P6 cerebellum, and P0 cerebral cortex of mice with the indicated genotypes. Cultured cells were stained for GFP (green), cilia markers (ACIII or ARL13B, red), biotin (fluorophore-conjugated streptavidin, white) and DNA (blue). Individual channels from the boxed regions are displayed to the right of each image in the following order (top to bottom): GFP, cilia marker, biotin and merge of the three channels. Scale bars: 5 µm (main) and 2.5 µm (insets). Scatter plots depict the fluorescence intensity of the ciliary biotin signal (relative to WT cilia, RFU: relative fluorescence unit) for cortical neurons **(C)**, GNPs **(E)**, and ependymal cells **(G)**. The thick and dotted lines indicate the median and interquartile range respectively. Number of cilia analyzed *N* = 110-159 from 3 independent experiments. Statistical significance was determined by one-way ANOVA with post-hoc Bonferroni test (****, *p* < 0.0001).

Primary cultures from diverse CNS cell types –cortical neurons, GNPs, and ependymal cells– revealed robust cilia-specific biotinylation only in cells from *Arl13b^+/TurboID^* mice (**Figs. 1B-G**). *In vivo*, prominent biotin signals were detected in cilia in the cerebral cortex and hippocampus of *Arl13b^+/TurboID^* mice at 1 and 3 months of age (**Figs. 2A and S2A-C**). Given that ciliary ARL13B levels are much lower in adult neurons than in glial cells *in vivo*, we expected ciliary biotinylation to be relatively weak in neuronal cilia in the brain of adult mice. Instead, the intensity of ciliary biotin signals was comparable between cilia marked by ARL13B (glial cells) or adenylate cyclase III (ACIII, neurons) in adult mouse cortex (**Fig. 2A-B**). As the levels of ciliary biotin were similar in cortical neurons from heterozygous and homozygous *Arl13b-TurboID* mice (**Fig. S2D-F**), we conclude that the modest levels of ARL13B-TurboID detected in neuronal cilia (**S2A-C**) are sufficient to carry out near-saturating levels of cilia-specific biotinylation.

**Figure 2.**
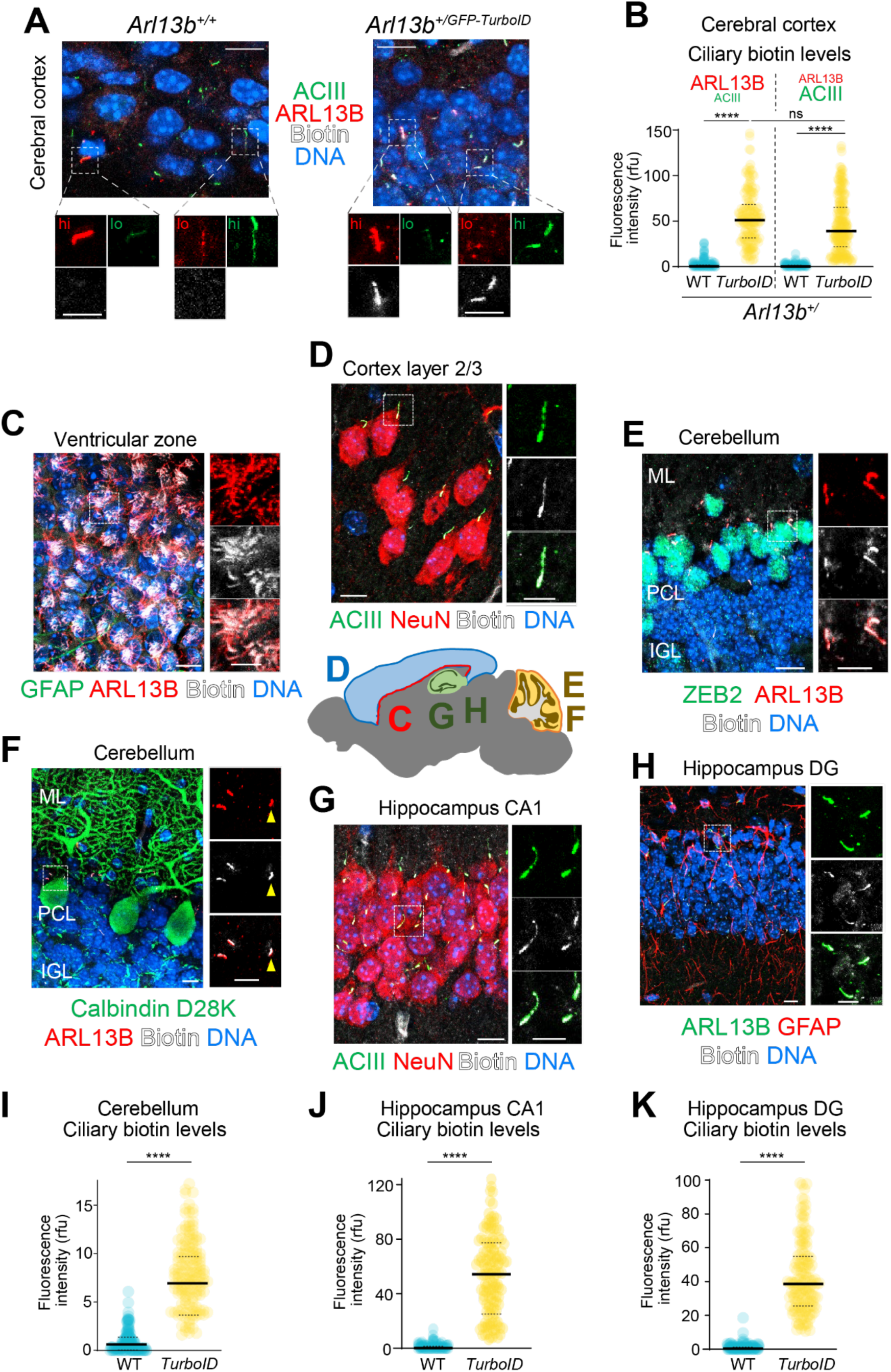
*In cilio* biotinylation by ARL13B-TurboID unveils a universal cilia marker. **A-B.** Robust ciliary biotinylation in *Arl13b^+/GFP-TurboID^* mice is independent of high ciliary levels of ARL13B. Following biotin injection for 3 days, sections of P21 cerebral cortex from WT (*Arl13b^+/+^*) and *Arl13b^+/GFP-TurboID^* mice were stained for the cilia markers ACIII (green) and ARL13B (red), for biotin (fluorophore-conjugated streptavidin, white) and for DNA (blue). Representative cilia with either high (hi) levels of ARL13B and low (lo) levels of ACIII or vice versa are boxed in each panel. Insets display individual channels from the boxed regions in the following order: ARL13B (top left), ACIII (top right) and biotin (bottom). Scale bar: 10 µm (main) and 5 µm (insets). Scatter plots (**B**) show the ciliary biotin levels relative to the levels in ARL13B^hi^/ACIII^low^ cilia in WT cerebral cortex. **C-K**. Cilium-specific biotinylation by ARL13B-TurboID in various brain regions after 3 days of biotin administration. Ventricular zone **(C)**, layer 2/3 of the cerebral cortex **(D)**, cerebellum **(E, F)**, as well as Cornu Ammonis 1 (CA1) **(G)** and dentate gyrus (DG, **H**) of hippocampus at P21 were stained with cilia markers (ACIII or ARL13B with indicated color), biotinylated proteins (fluorophore-conjugated streptavidin, white), DNA (blue), and cell-type specific markers: post-mitotic neurons (NeuN, red in **D** and **G**), glial cells (GFAP, green in **C** and red in **H**), Bergmann glial cells (ZEB2, green in **E**), and Purkinje neurons (Calbindin D28K, green in **F**). Arrowheads in 2F specify the cilium of Purkinje neurons. Abbreviations: ML: molecular layer; PCL: Purkinje cell layer; IGL: internal granular layer of the cerebellar cortex. Scatter plots show the fluorescence intensity of the ciliary biotin signal (relative to WT cilia) in the cerebellum **(I)**, hippocampus CA1 **(J)**, and hippocampal DG **(K)**. Scale bar: 10 µm (main) and 5 µm (insets). RFU: relative fluorescence unit. Number of cilia analyzed: *N* = 124-239 **(B)**, 108-178 **(I, J, K)** from 3 mice. Thick and dotted lines represent the median and interquartile region in all scatter plots. Statistical significance was determined by Two-way ANOVA with post-hoc Bonferroni test in **(B)**; Mann-Whitney U test in **(I-K)** (****, *p* < 0.0001; ns: not significant).

Encouraged by these results, we examined the breadth of ARL13B-TurboID biotinylation across different mouse tissues. Robust biotin signals were detected in motile cilia of ependymal cells lining the ventricular zone (**Fig. 2C**) and in primary cilia of cortical pyramidal neurons (**Fig. 2D**), Bergmann glial cells (**Fig. 2E**), Purkinje neurons (**Fig. 2F**), hippocampal principal neurons (**Fig. 2G**), and glial cells of the dentate gyrus (**Fig. 2H**). Strong ciliary biotin signals were also observed in the distal convoluted tubules of the kidney and the taste buds of the tongue (**Fig. S3A-B**) and ciliary biotin signals in all tested tissues were specifically detected in *Arl13b-TurboID* mice (**Figs. 2I-K and S3C-D**). These observations highlight the utility of the *Arl13b-TurboID* mouse as a universal marker of cilia, surpassing conventional markers such as ARL13B, ACIII, and acetylated tubulin, which label cilia only in specific cell types.

### Proteomic profiling Reveals Tissue-Specific Specialization of Ciliary Proteomes

The biochemical isolation of biotinylated proteins revealed that many proteins were selectively recovered from *Arl13b-TurboID* cerebral cortices, where their levels were enhanced by biotin injection (**Fig. S4A**). Importantly, the untagged ARL13B protein, an endogenous ciliary component, was recovered from *Arl13b-TurboID* samples but not from *Arl13b[V358A]-TurboID* and wild-type samples, demonstrating that only ARL13B-TurboID can biotinylate ciliary proteins (**Fig. S4B**).

To compare ciliary proteomes across different organs, we conducted quantitative proteomic profiling of biotinylated proteins isolated from triplicate or quadruplicate hippocampus and kidney lysates from *Arl13b^+/TurboID^* and *Arl13b^+/+^* mice. Tandem Mass Tag (TMT) analysis quantified nearly 1200 proteins, and hierarchical clustering identified 100 proteins specifically enriched in *Arl13b^+/TurboID^* samples (**Fig. S4C, Table S1**). Notably, less than a third of these proteins were shared between hippocampal and renal ciliary proteomes, including ARL13B, INPP5E^41^, and NUMBL^42^. Meanwhile, more than a third were present exclusively in the kidney-enriched cluster –including well-known ciliary components such as MYO6^34,43^, TULP3^44^, and IFT74^45^– and another third were only found in the hippocampus-enriched cluster. These findings highlight an unexpected diversity of ciliary proteomes across tissues, contrasting the well-characterized renal cilia proteome^34,35,46^ with the poorly understood neuronal cilia proteome.

### *In situ* proteomics uncovers known and novel ciliary proteins in the brain

Our experience with cilia proteomics^35^ has emphasized that genetic controls (e.g. ablating cilia) are superior to conventional controls (e.g. delocalized enzyme) for identifying bona fide ciliary proteins. To this end, we combined the *Arl13b-TurboID* allele with homozygous *Ift88^flox^* alleles to genetically remove cilia in a Cre-dependent manner. A ubiquitous tamoxifen-activated Cre (*Ubc-cre/ERT2*) was used to delete all cilia in adult mice (TurboID/cilia) and we employed *Nex-cre*^47^ to selectively ablate cilia in pyramidal neurons (TurboID/cilia^neu^). Additionally, we combined *Arl13b-TurboID* with a *Bbs4* deletion to impair BBSome-dependent retrieval mechanisms, allowing accumulation of ciliary proteins^48^ (TurboID/Bbs, **Fig. 3A**). Efficient cilia ablation and ciliary biotinylation were confirmed via imaging cortical neurons and glial cells (**Fig. 3A-E and S5A-B**). Although cilia were shorter in *Bbs4^-/-^* cells (**Figs. S5A-B**), ciliary biotinylation was unaffected (**Fig. 3C & E**).

**Figure 3.**
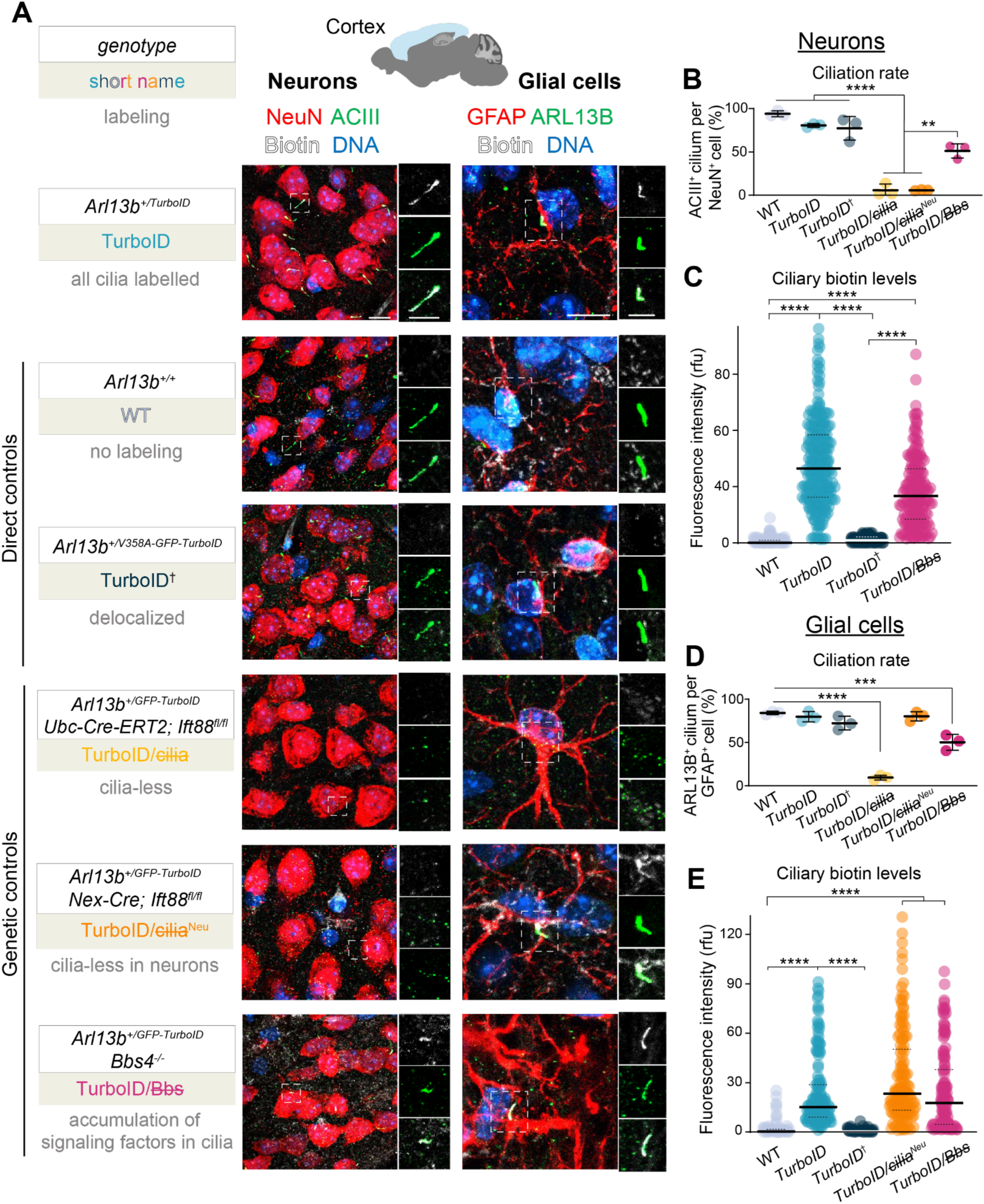
Genetic and direct controls for cilia proteomics. **A.** Ciliation and ciliary biotinylation in neurons and glial cells within the cerebral cortex across six genotypes. Following 3 days of biotin injection, sections of cerebral cortices from the indicated genotypes were stained with markers of cilia and neurons (ACIII, green; NeuN, red) or cilia and glial cells (ARL13B, green; GFAP, red) along with biotin (fluorophore-conjugated streptavidin, white), and DNA (blue). The tables on the left of each panel provide information in the following order (top to bottom): genotype, short name of the genotype, and effects of biotin labeling. Individual channels from boxed regions are enlarged to the right of each panel in the following order: biotin, cilia marker, and merge of these two channels. Scale bar: 10 µm (main) and 5 µm (insets). **B and D**. Quantitation of ciliation rate in cells. Plots represent the percentage of ACIII^+^ cilia among NeuN^+^ neurons **(B)** and of ARL13B^+^ cilia among GFAP^+^ glial cells **(D)** for the indicated genotypes. *N* = 3 animals per genotype. Lines mark the mean and standard deviation. **C and E**. Quantitation of biotin levels in cilia. Scatter plots depict the fluorescence intensity of the ciliary biotin signal (relative to WT cilia, RFU: relative fluorescence unit) for neurons **(C)**, and glial cells **(E)**. Thick and dotted lines represent the median and interquartile range, respectively. Number of cilia analyzed: *N* = 105-227 **(B-E)** from 3 brains per genotype. Statistical significance was determined by one-way ANOVA with post-hoc Bonferroni test (**, *p* < 0.01; ***, *p* < 0.001; ****, *p* < 0.0001). See **Figs. S5A-B** for additional measurements.

To broaden proteomic coverage, we implemented two complementary purification strategies: one capturing all proteins via detergent extraction and another enriching membrane-associated proteins using wheat germ agglutinin (WGA) beads (**Fig. 4A**). Brain samples from four to five experimental conditions in triplicate or quadruplicate were analyzed using quantitative mass spectrometry, and clustering analysis identified groups of proteins enriched in brain cilia (TurboID > TurboID^†^ ∼ WT, **Fig. 4B**, or TurboID > TurboID/cilia ∼ WT, **Fig. 4C**) or neuronal cilia specifically (TurboID/Bbs > TurboID > TurboID/cilia^Neu^ > WT, **Fig. 4D**). Western blot analyses confirmed the effective purification of biotinylated proteins (**Fig. 4B-D**).

**Figure 4.**
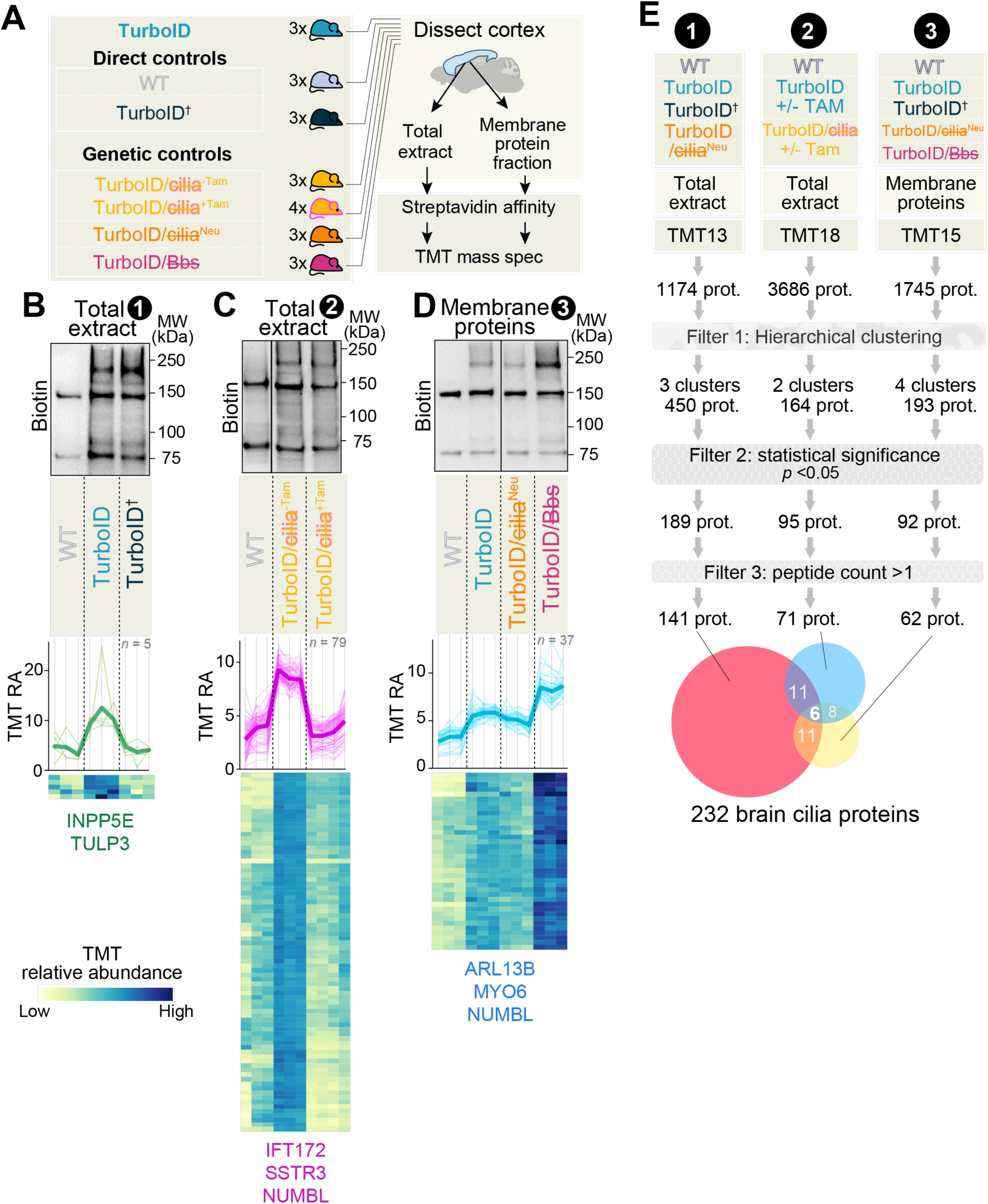
Proteomic profiling identifies candidate cilia proteins in the cerebral cortex. **A.** Diagram of the experimental workflow for cilia proteomics. Genotype abbreviations: TurboID: *Arl13b^+/GFP- TurboID^*. WT: *Arl13b^+/+^*. TurboID†: *Arl13b^+/V358A-GFP-TurboID^*. TurboID/cilia: *Arl13b^+/GFP-TurboID^; Ubc-cre/ERT2; Ift88^fl/fl^*. TurboID/cilia^Neu^: *Arl13b^+/GFP-TurboID^; Nex-cre; Ift88^fl/fl^*. TurboID/Bbs: *Arl13b^+/GFP-TurboID^; Bbs4^-/-^*. Tam: Tamoxifen. Cerebral cortex lysates (total extracts or membrane-associated fractions) from indicated genotypes were subjected to streptavidin purification followed by tandem mass tag (TMT) mass spectrometry. The number of replicates used in each TMT-MS experiment are indicated next to the mouse genotype. **B-D.** Representative clusters of candidate ciliary proteins in three independent TMT-MS experiments. Total extracts (**B-C**) and membrane fractions (**D**) from dissected cortices were purified on streptavidin beads and analyzed by immunoblotting (upper panels) or TMT-MS (middle and lower panels). Upper panels: 40 equivalents of eluates were loaded in the lysate lanes. Middle panels: line graphs depict the relative abundance (RA) of a given protein, calculated via dividing the TMT signals in each sample by the sum of the TMT signals across all samples. Thick lines show the average RA for each protein cluster, and thin lines show individual proteins. Lower panels: heatmaps display the RA of each protein across the genotypes. The color scheme at the bottom left indicates the RA scale. See **Figs. S5C-I** for additional clusters. **E.** Analytical pipeline to identify candidate brain cilia proteins. Three independent TMT-MS experiments (see **Table S2**) were subjected to hierarchical clustering analysis. Clusters with profiles expected for cilia proteins were pooled (filter 1), hits were filtered for statistical significance (filter 2: *p* < 0.05 for TurboID vs. WT) and peptide numbers (filter 3: peptide count > 1). A Venn diagram of overlapping proteins in experiment 1 (red), 2 (blue), and 3 (yellow) from the final tier. The numbers of proteins shared among datasets are shown.

In TMT experiment ❶ (**Table S2**), hierarchical clustering of TurboID vs. WT samples revealed a cluster enriched in known ciliary proteins, such as ß-tubulin4B (TUBB4B)^49^ (**Fig. S5C**). Another cluster, matching the neuronal cilia profile TurboID > TurboID/cilia^Neu^ > WT, included the known cilia proteins myosin VI (MYO6) and SNAP25^50^ (**Fig. S5D**). Surprisingly, only five proteins – including tubby-like protein 3 (TULP3) and inositol polyphosphate 5-phosphatase E (INPP5E)– matched the profile TurboID > TurboID^†^ ∼ WT (**Fig. 4B**), suggesting that only a few cilia proteins are exclusively localized to cilia in brain cells.

In TMT experiment ❷ (**Table S2**), tamoxifen (Tam) was added to half of the TurboID and TurboID/cilia mice to control for the effects of this drug. A cluster enriched in TurboID vs. WT samples included ARL13B, MYO6, and INPP5E (**Fig. S5E**). Another cluster with the profile TurboID^-Tam^ ∼ TurboID^+Tam^ and TurboID/cilia^-Tam^ > TurboID/cilia^+Tam^ ∼ WT identified brain cilia proteins and included IFT172^51^, NUMBL, and the ciliary GPCR somatostatin receptor 3 (SSTR3) (**Figs. 4C and S5F**). The detection of SSTR3 in our dataset highlight the sensitivity of the method, given the low abundance of GPCRs in neuronal cilia (<1000 molecules per cilia)^40^.

In TMT experiment ❸ (**Table S2**), membrane-associated proteins were purified from cortical lysates of WT, TurboID, TurboID^†^, TurboID/cilia^neu^, and TurboID/Bbs mice. Three clusters of candidate neuronal cilia proteins (TurboID/Bbs > TurboID > TurboID/cilia^neu^ > WT) included the known ciliary proteins ARL13B, INPP5E, SNAP25, MYO6, and NUMBL, and the metabotropic GPCRs GABA_B1_^52^ and the glycine receptor GPR158^53^ (**Fig. S5G-H**). Among these, 37 proteins showed a pronounced enrichment in retrieval mutants, underscoring the dynamic shuttling of ciliary proteins (**Figs. 4D and S5H**). An additional cluster enriched in TurboID vs. WT included IFT172 and BBS5^40^ (**Fig. S5I**).

Collectively, these proteomic analyses across different genotypes revealed the diversity of known and novel ciliary proteins, offering new insights into neuronal cilia biology.

### Synaptic proteins are selectively enriched in brain cilia

A high-confidence brain cilia proteome was refined to 232 candidate brain cilia proteins by filtering for statistical significance and peptide counts (**Fig. 4E, Table S2**). Remarkably, less than 18% of these proteins are found in the published ciliary proteomes of cultured kidney cells^34,35,46,54^ (**Table S2**), further underscoring the specialization of brain cilia. Gene ontology analyses revealed a remarkable enrichment of synaptic proteins in neuronal cilia (**Fig. S6, Table S3**), including the GluN1 subunit of the ionotropic NMDA receptor, the GABA_B1_ subunit of the metabotropic GABA receptor, several Na^+^/K^+^ ATPase subunits, and the voltage-gated sodium channel Na_V_1.1 (**Fig. 5A**). Furthermore, the brain cilia proteome also included multiple adhesion molecules, such as neuron cell adhesion molecule 1 (NCAM1) and the tight junction protein ZO-1 (**Fig. 5A**). These findings challenge the traditional view that synaptic signaling in neurons is confined to dendrites and suggest that neuronal cilia may function as specialized extrasynaptic devices.

**Figure 5.**
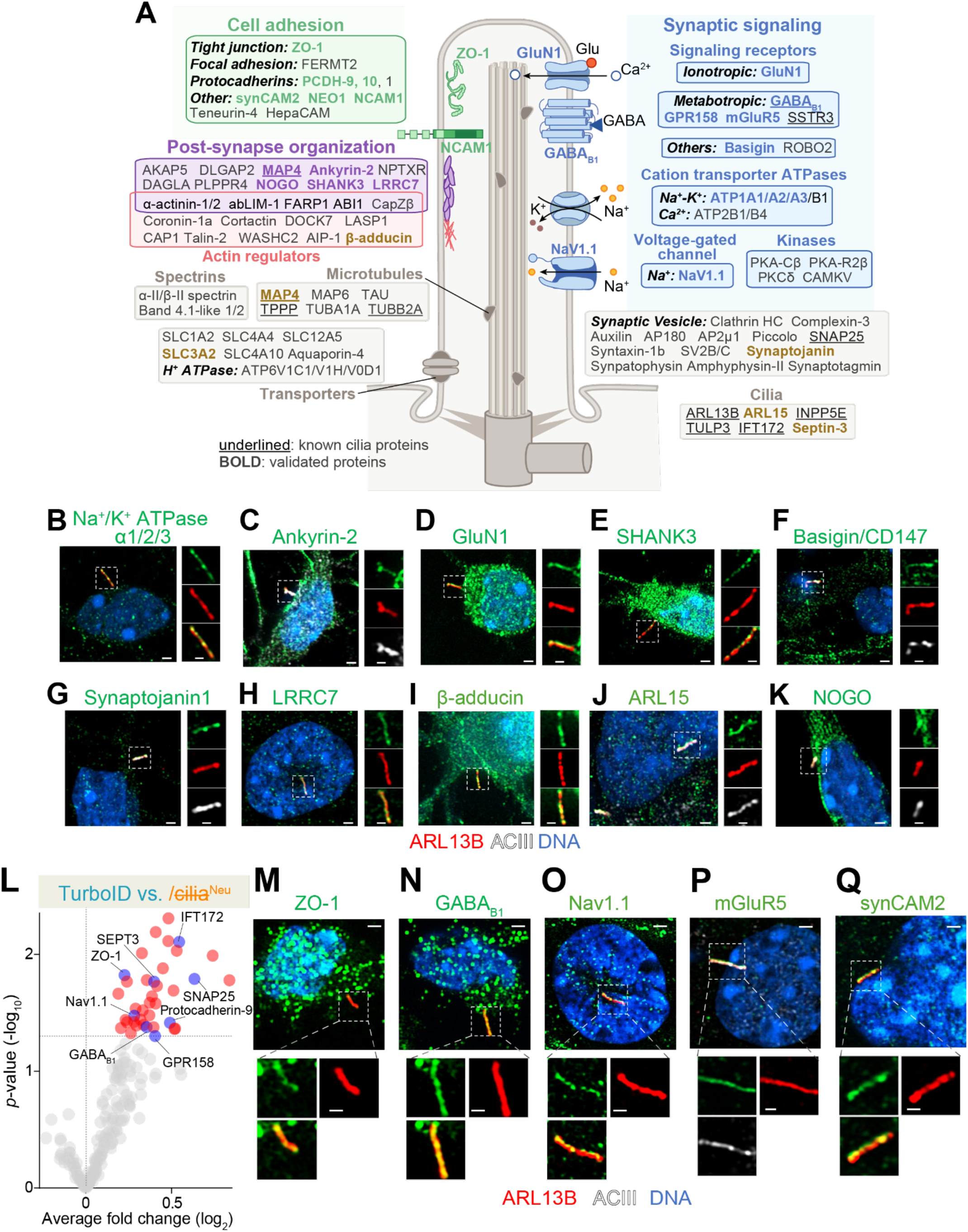
Overview of the brain and neuronal cilia proteome. **A.** Overview of the brain cilia proteome with functional groupings in synaptic reception (blue), cell adhesion (green), post-synaptic organization (magenta), or other functions (grey). Candidates validated by immunostaining in this study are **bolded**. **B-K**. Validation of the brain cilia proteome. Cultured cortical neurons (DIV8) from E15.5 cerebral cortex were stained for candidate cilia proteins (green), ciliary markers (ARL13B, red; ACIII, white), and DNA (blue). The antibody used in panel **B** recognizes ATP1A1, A2 and A3. Individual channels from boxed regions are enlarged to the right of each panel with the following order (top to bottom): candidate protein, ARL13B, and either merged channels (for **B, D, E, H, and I**) or ACIII (for **C, F, G, J, and K**). Images captured with Airyscan2. Scale bar: 1 μm (main) and 500 nm (insets). **L-Q**. Validation of the neuron-enriched cilia proteome. **L**. Volcano plots comparing TurboID (*Arl13b^+/GFP-TurboID^*) vs. TurboID/cilia^Neu^ (*Arl13b^+/GFP-TurboID^; Nex-cre; Ift88^fl/fl^*) TMT signals amongst 186 Tier 1 proteins identified in TMT experiments 1 and 3. Statistical *p* values are plotted against TMT relative abundance ratios in each comparison. Proteins enriched in TurboID samples are highlighted in red. Known cilia proteins and validated proteins are labeled in blue, and all other proteins are shown in gray (see **Table S4** for dataset). **M-Q**. Cultured cortical neurons (DIV8) from E15.5 cerebral cortex were stained for candidate cilia proteins (green), ciliary markers (ARL13B, red; ACIII, white), and DNA (blue). Individual channels from boxed regions are enlarged to the bottom of each panel with the following order: candidate protein (top left), ARL13B (top right), and either merged channels (for **M, N, O, and Q**) or ACIII (**P**). Images captured with Airyscan2. Scale bar: 1 μm (main) and 500 nm (insets).

Immunostaining cultured cortical neurons using well-validated antibodies (**see Methods**) validated several candidates localized to neuronal cilia. The Na^+^/K^+^ ATPase and its scaffolding protein ankyrin-B were detected in cilia, supporting their role in maintaining ionic homeostasis (**Fig. 5B-C**). Similarly, the NMDA receptor subunit GluN1 and its scaffolding protein SHANK3 were detected in neuronal cilia (**Fig. 5D-E**), highlighting their potential involvement in neurotransmitter sensing. Additional validated proteins include the signaling receptor basigin, the phosphoinositide phosphatase synaptojanin1, the postsynaptic scaffold protein LRRC7, and the actin and spectrin regulator β-adducin (**Fig. 5F-I**). Notably, the ARF-like GTPase ARL15 was also detected in neuronal cilia (**Fig. 5J**), representing the fourth member of this family (after ARL3, ARL6, and ARL13B) to function within cilia. Finally, the regulator of neurite outgrowth NOGO was found in neuronal cilia (**Fig. 5K**). Collectively, these results demonstrate that neuronal cilia harbor several postsynaptic proteins, including neurotransmitter receptors, scaffolding proteins, and signaling regulators.

Among the brain cilia proteins, a subset of candidate pyramidal neuron cilia proteins was identified by comparing the TurboID and TurboID/cilia^Neu^ datasets (**Fig. 5L, Table S4**). These candidates include protocadherin-9, GPR158, GABA_B1_, Na_V_1.1, ZO-1, septin-3 (SEPT3), IFT172, and SNAP25. Immunostaining of cultured cortical neurons confirmed the ciliary localization of ZO-1 (**Fig. 5M**), GABA_B1_ (**Fig. 5N**) and Na_V_1.1 (**Fig. 5O**). These findings suggest that pyramidal neuron cilia may mediate cell adhesion, synaptic signal reception, and neuronal excitability.

Mapping the brain cilia proteome onto single-cell RNA sequencing data^55^ (**Fig. S7, Table S4**) revealed that some proteins – e.g. NCAM1– appeared broadly expressed in most ciliated cell types, while others –e.g. the metabolic glutamate receptor mGluR5, synaptic cell adhesion molecule 2 (SynCAM2), and SNAP25– were highly expressed in glutamatergic and GABAergic neurons (**Fig. S7**). Staining of cultured cortical neurons validated the presence of mGluR5 and SynCAM2 in neuronal cilia (**Fig. 5P, Q**).

Altogether, these results underscore a significant convergence between the neuronal cilia and synaptic proteomes, suggesting that neuronal cilia exploit synaptic machinery to fulfill their functions.

### Many brain cilia proteins are not exclusively ciliary

Building on the notion that neuronal cilia may co-opt proteins from other cellular compartment, only eight proteins –including the known ciliary proteins IFT172 and microtubule-associated protein 4 (MAP4)^56^, along with the novel ciliary candidate septin-3 and μ-crystallin (CRYM)– were significantly enriched in TurboID samples compared to TurboID^†^ controls (**Fig. 6A, Table S5**). In parallel, comparing TurboID/cilia^-Tam^ with TurboID/cilia^+Tam^ samples revealed 16 proteins significantly enriched in TurboID/cilia^-Tam^ eluates (**Fig. 6B, Table S5**). This group comprised canonical ciliary components (SSTR3, IFT172, INPP5E), as well as novel candidates such as septin-3, SLC3A2 (the large subunit of neutral amino acid transporters). These subsets of proteins were classified as Tier 1* due to their robust ciliary enrichment.

**Figure 6.**
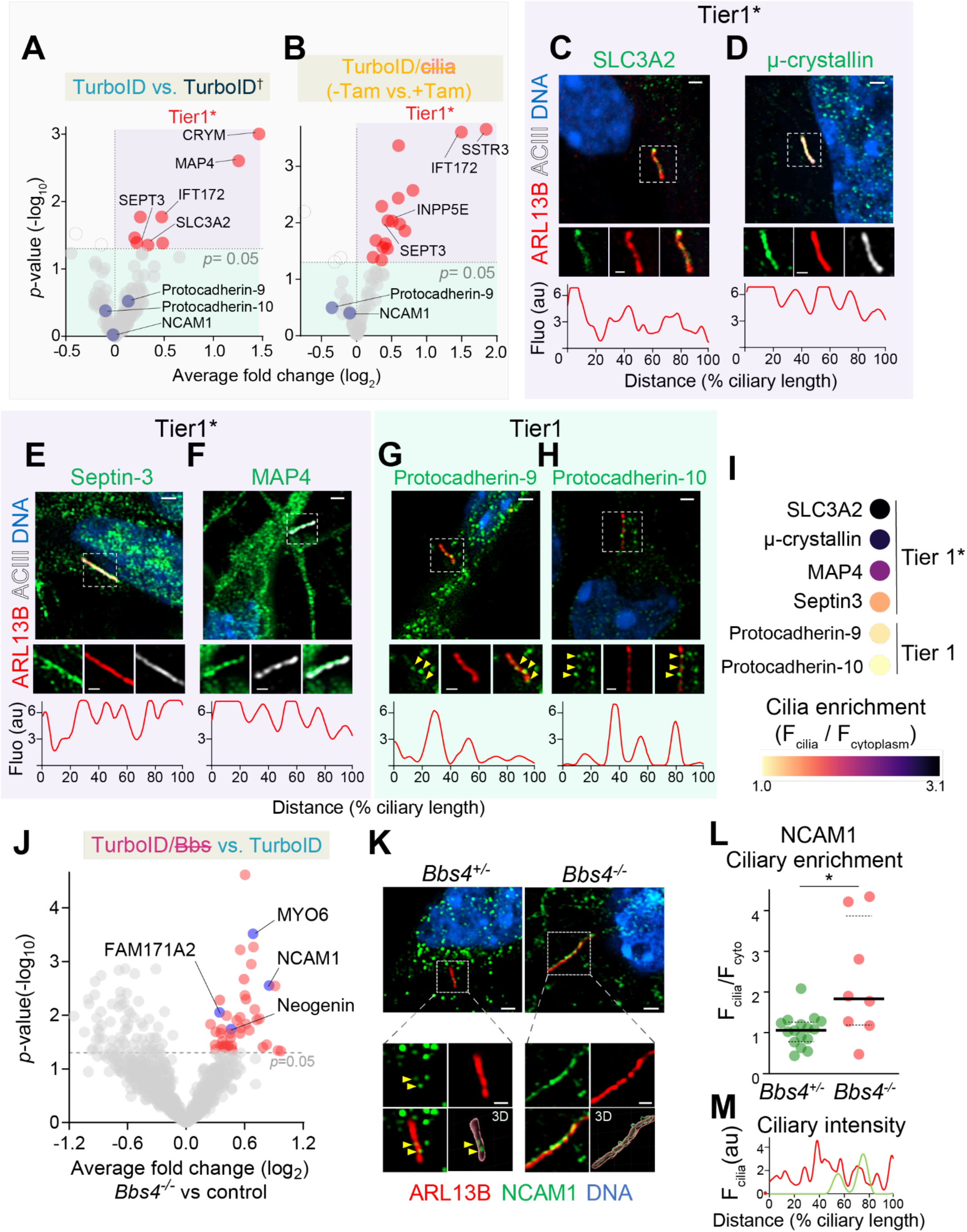
Novel ciliary proteins show differential ciliary enrichment. **A, B.** Volcano plots comparing (**A**) TurboID (*Arl13b^+/GFP-TurboID^*) vs. TurboID^†^ (*Arl13b^+/V358A-GFP-TurboID^*) TMT signals amongst 186 Tier 1 proteins identified in TMT experiments 1 and 3. (**B**) TurboID/cilia^-Tam^ vs. TurboID/cilia^+Tam^ (*Arl13b^+/GFP-TurboID^; Ubc-cre/ERT2; Ift88^fl/fl^* ± tamoxifen) with 71 Tier 1 proteins identified in TMT experiment 2. Statistical *p* values are plotted against TMT relative abundance ratios in each comparison. Proteins significantly enriched in TurboID or TurboID/cilia^-Tam^ samples are highlighted in red and classified as Tier 1* whereas all other Tier 1 proteins are shown in gray and validated proteins shown in blue. Proteins significantly de-enriched are displayed with a dashed outline; these proteins were removed from the brain cilia proteome (see **Table S5** for dataset). **C-H**. Validation of candidate Tier 1* and Tier 1 protein localization to neuronal cilia by Airyscan2 super-resolution imaging. Cultured cortical neurons (DIV8) were stained for candidate proteins (green), ciliary markers (ARL13B or ACIII, as indicated in red or white channels), and DNA (blue). The boxed regions are enlarged to the bottom of each panel with the following order (left to right): candidate protein, cilia marker, and merged channels (for **C**, **F**, **G**, **H**) or ACIII (for **D**, **E**). Arrow heads in insets of **G** and **H** indicate candidate proteins on cilia. Scale bars: 1 µm (main) and 500 nm (insets). **I**. Dot plots depict the cilia enrichment of 6 candidate proteins selected from Tier 1* and 1. Cilia enrichment is defined by the ratio of fluorescent signals in cilia to the rest of cells of a given protein. Each dot is color-coded by scaled average ciliary enrichment (ranging from 1.0 to 3.1). *N* = 15-68 cilia were collected for each protein from 3 independent experiments. See **Fig. S8A** for details. **J**. Proteins enriched in cilia of *Bbs4^-/-^* versus control cortices. Volcano plots comparing the proteomes of *Bbs4^-/-^* (*Arl13b^+/GFP-TurboID^; Bbs4^-/-^*) vs. control (*Arl13b^+/GFP-TurboID^*) cortices with 1745 proteins identified in WGA, Tier3. Statistical *p* values are plotted against TMT ratios. Proteins enriched in *Bbs4^-/-^* cilia are highlighted in red, known cilia proteins and validated proteins are labeled in blue, and all other proteins are shown in gray (see **Table S6** for dataset). **K-M**. The Tier 1 protein, NCAM1 is enriched in *Bbs4^-/-^* neuronal cilia. **K**. Cultured cortical neurons (DIV8) from E15.5 *Bbs4^+/-^* (left) or *Bbs4^-/-^* (right) cerebral cortices were stained for NCAM1 (green), a ciliary marker (ARL13B, red), and DNA (blue). The boxed regions are enlarged to the bottom of each panel with the following order: NCAM1 (top left), ARL13B (top right), merged channels (bottom left), and 3D reconstruction (Bottom right). Arrow heads in insets indicate the candidate protein in cilia. Scale bars: 1 µm (main) and 500 nm (insets). **L**. Plots show the ratio of ciliary to cytoplasmic fluorescence for NCAM1 in individual cells. Thick and dotted lines represent the median and interquartile region. *N*=8-16 cilia scanned by Airyscan2 super-resolution module. Statistical significance was determined by the Mann-Whitney U test (*, *p* < 0.05). **M**. Linescans reveal the fluorescence of NCAM1 (arbitrary unit, au) through the *Bbs4^+/-^* (green) and *Bbs4^-/-^* (red) cilia. The length of cilium is normalized to 100%.

Immunostaining of cultured cortical neurons confirmed that SLC3A2, μ-crystallin, septin-3, and MAP4 are highly concentrated in cilia (**Fig. 6C-F**). While Tier 1* proteins exhibited continuous staining patterns along cilia, other Tier 1 proteins, such as protocadherin-9 and -10, showed punctate localization (**Fig. 6G-H**). Quantifying enrichment ratios revealed that Tier 1* proteins are markedly concentrated in cilia, whereas other Tier 1 proteins demonstrated more modest enrichment (**Figs. 6I and S8A**). Concordantly, mapping the brain cilia proteome onto an organelle atlas^57^ revealed that most brain cilia proteins also localize to other compartments, chiefly the plasma membrane, cytosol and cortical actin (**Fig. S8B, Table S5**).

Furthermore, retrieval-deficient *Bbs4^-/-^* neurons showed significant accumulation of proteins such as NCAM1, FAM171A2, and neogenin (**Fig. 6J, Table S6**), which exhibited punctate patterns in control cilia and a more continuous distribution in *Bbs4^-/-^* cilia (**Figs. 6K-M and S8C-H**). This accumulation was not associated with global increases in protein levels (**Fig. S8I-J**), suggesting dynamic shuttling of proteins into and out of cilia mediated by the retrieval machinery save for specific nanodomains within the cilium.

### Visualization of ciliary proteins within the brain ultrastructure reveals close association of ciliary NMDA receptors with synapses

To explore the spatial relationship between neuronal cilia proteins and the brain architecture *in vivo*, we employed <u>pan</u>-<u>ex</u>pansion <u>m</u>icroscopy of tissues (pan-ExM-t)^58^, achieving over 16-fold linear expansion of brain samples (**Fig. S9A-B**) and bringing resolution below 20 nm. Fluorescent NHS-esters (‘pan’-stains) enabled near-ultrastructural visualization of brain architecture, including cilia and basal bodies (**Fig. 7A-B**). Immunostaining revealed the ciliary membrane protein ACIII decorating a cylindrical pattern, with axonemal microtubules and basal body stained by NHS-ester (**Fig. 7B**). Excitatory synapses were readily identified as parallel asymmetric structure in NHS ester staining, and the location of the post-synaptic compartment^59^ and perisynaptic region^60^ were confirmed by NMDAR and protocadherin-9 staining, respectively (**Fig. S9C**). Consistent with prior EM studies^28^, synapses were frequently seen within the vicinity of neuronal cilia in pan-ExM-t (**Fig. 7C, Video S1**).

**Figure 7.**
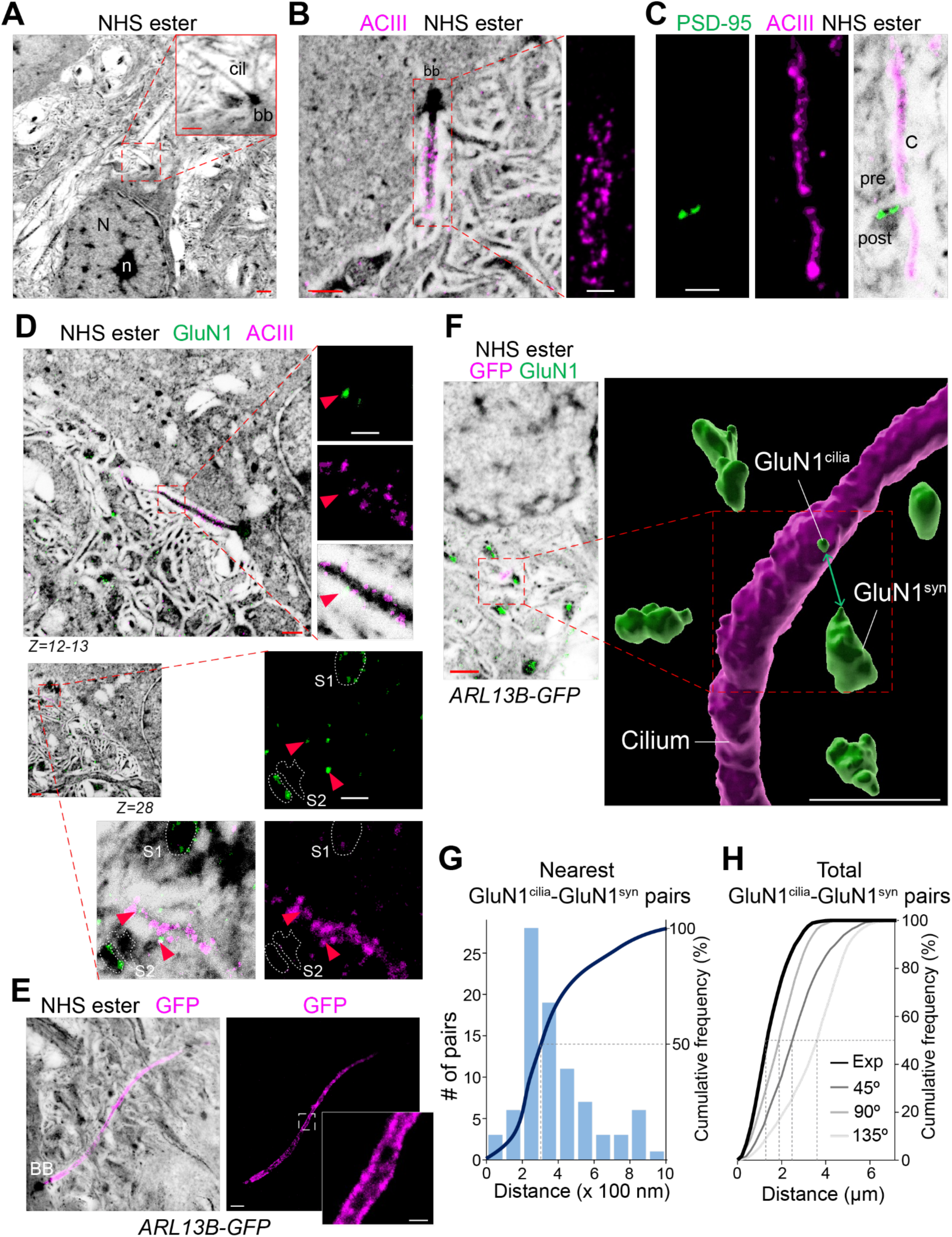
Ciliary NMDA receptors are positioned near neighboring synapses. **A.** Visualization of neuronal cilium via pan-stained expansion microscopy of tissues (pan-ExM-t). Cerebral cortices from P60 mice were expanded 16.2-fold and stained with fluorescent NHS-ester (black). The red boxed region is enlarged at the upper right of this panel. N: nucleus, n: nucleolus, cil: cilium, bb: basal body. Scale bar: 1 μm (main) and 500 nm (inset). **B.** ACIII distribution on the ciliary membrane in pan-ExM-t. Expanded cerebral cortices from P60 mice were stained for ACIII (magenta) and with fluorescent NHS-ester (black). ACIII signals in the boxed region are enlarged to the right. Scale bars: 500 nm (main) and 250 nm (inset). Representative images from three independent trials with over 25 cilia imaged. **C.** Visualization of neuronal cilia and synapses in pan-ExM-t. P60 cerebral cortices were expanded and stained for the postsynaptic marker PSD-95 (green), the ciliary membrane marker ACIII (magenta) and NHS ester (black). Individual channels are shown from left to right (green, magenta, and merged). C: cilium, pre: pre-synapse, post: post-synapse. Scale bar: 250 nm. Representative images from three independent trials with over 10 cilia imaged. See also **Video S1**. **D.** GluN1 localization on the ciliary membrane adjacent to synapses. Expanded P60 cerebral cortices were stained for ACIII (magenta), GluN1 (green) with NHS-ester (black). Boxed regions are enlarged, showing GluN1, ACIII, and merged channels with NHS-ester from top to bottom (optical section: Z = 12-13) and top, bottom right, and bottom left (z=28). Red arrowheads: ciliary GluN1 signals near two synapses (S1 and S2). Scale bars: 500 nm (main) and 200 nm (insets). Representative images from three independent trials with over 15 imaged. See also **Video S3**. **E.** Expanded P45 cerebral cortex from *ARL13B-GFP^tg^* mice stained for GFP (magenta) and with NHS-ester (black). Inset in the GFP channel shows the enlarged image from the boxed region. Representative image from three independent trials with over 30 cilia imaged. Scale bar: 500 nm (main), 100 nm (insets). **F.** Left: expanded cerebral cortices from P45 *ARL13B-GFP^tg^* mice were stained for GFP (magenta), GluN1 (green) and with NHS-ester (black). Right: 3D rendering of cilia (from the GFP channel) and GluN1 structures, either synapses (GluN1^syn^) or ciliary nanodomains (GluN1^cilia^) from 180 Z sections. The image on the left is a sum of intensity projection from Z= 69-70. The green double arrow marks a representative pair of GluN1 structures on cilia and on synapses. Scale bars: 500 nm (3D rendering and original image). See also **Video S4**. **G.** Histogram depicting the shortest distance in 3D between GluN1^cilia^ spots and the nearest GluN1^syn^ structure. The blue bars display the number of GluN1^cilia^-GluN1^syn^ pairs (as shown by the green line in **F**) for a given range of distances. The black line shows the cumulative frequency of distances for the nearest GluN1^cilia^-GluN1^syn^ pairs, and the dashed line marks the median at 297 nm. *N* = 9 cilia from 3 independent trials were segmented and quantified. **H.** Cumulative frequencies of distances between GluN1^cilia^ spots and GluN1^syn^ structures in a 3D volume. The solid lines represent the cumulative frequencies of distances for total GluN1^cilia^-GluN1^syn^ pairs. The black line represents the experimental values. To test whether the observed distribution is non-random, cilia were rotated by 45°, 90°, and 135° (greys) before conducting the analysis. The dashed lines mark the medians at 1.43 μm (experimental), 2.60 μm (45°), 1.97 μm (90°), 3.70 μm (135°). *N* = 9 cilia from 3 independent trials were segmented and more than 3200 GluN1^cilia^-GluN1^syn^ pairs were quantified. All scale bars are corrected for the expansion factor. All micrographs shown are maximal intensity projections of the Z sections spanning a neuronal cilium, except for the bottom panels in **D** where a single Z section is shown.

The unprecedented resolution of pan-ExM-t enabled us to unequivocally determine the specificity of discrete signals of signaling receptors decorating cilia as their position matched that of ciliary membrane markers such as ACIII or ARL13B. For instance, co-staining for the metabotropic glycine receptor GPR158^53^ and ACIII revealed discrete foci on the membrane of neuronal cilia *in vivo* (**Video S2**), extending a recent report of its ciliary localization in cultured neurons^61^.

Equipped with a powerful tool to precisely locate ciliary signaling receptors and synapses inside the brain architecture, we zeroed in on the NMDA receptor subunit GluN1 as it was at the center of a major node of glutamatergic postsynaptic proteins in protein association networks of the brain cilia proteome (**Fig. S9D**). Interestingly, discrete GluN1 foci specifically decorated the ciliary membrane cylinder marked by ACIII *in vivo* (**Fig. 7D top, Video S3**). Notably, the discrete GluN1 signals on ciliary membranes were consistently positioned in close proximity to neighboring GluN1-positive glutamatergic synapses (**Fig. 7D bottom**), suggesting an organized spatial relationship between ciliary NMDA receptors and nearby synaptic structures.

To rigorously test whether ciliary GluN1 molecules are non-randomly positioned relative to synapses, we stained cilia in *ARL13B-GFP* cortical sections (**Fig. 7E**) and performed automated 3D segmentation of neuronal cilia and synapses (**Fig. 7F, Video S4**). Quantifying the shortest distances between ciliary GluN1 molecules and the nearest neighboring synapses revealed a median distance just under 300 nm (**Fig. 7G**), very similar to the shortest distance between the ciliary membrane and neighboring synapses (**Fig. S9E**). Further emphasizing the close relationship between ciliary GluN1 nanodomains and synapses, this value is in close agreement with volume EM-based measurements of cilia-synapse distances in the mouse visual cortex^28^. Randomizing cilia orientation disrupted this spatial association (**Fig. 7H**), demonstrating that ciliary NMDA receptors are non-stochastically positioned near synapses.

In conclusion, the mapping of synaptic structures and proteins on neuronal cilia in brain volumes at near-ultrastructural resolution by pan-ExM-t extends previous electron microscopy-based studies^27–29^ and reveals nanodomains on neuronal cilia strategically located to monitor activity from nearby glutamatergic synapses.

## DISCUSSION

### Neuronal Cilia as Extrasynaptic Signaling Platforms

Neuronal cilia have long been implicated in brain development^19^ and connectivity^62^, but their roles in mature neural circuits remain poorly understood. This study provides compelling evidence that neuronal cilia functions as specialized extrasynaptic signaling platforms, equipped with synaptic receptors, adhesion molecules, and regulators of neurite outgrowth (**Fig. 5A**). By profiling the brain cilia proteome and visualizing nanoscale protein organization, we demonstrate that neuronal cilia are positioned to transduce signals from neighboring synapses (**Fig. 7**), potentially influencing neuronal excitability and connectivity.

The discovery of synaptic proteins, most notably the NMDA receptor subunit GluN1 (**Fig. 5D**), scaffolding proteins such as SHANK3^63^ (**Fig. 5E**), and signaling molecules such as the metabotropic GABA receptor subunit GABA_B1_ (**Fig. 5N**), within cilia challenges the conventional view that synaptic signaling in neurons is confined to dendrites. Our expansion microscopy analysis revealed a discrete localization of GluN1 on the ciliary membrane near glutamatergic synapses (**Figs. 7D**, **F-H**), suggesting that neuronal cilia are equipped to integrate local synaptic activity. These findings extend previous electron microscopy studies that identified axo-ciliary synapses and synapse-cilia associations in the mouse and human brains^27–29^. The molecular framework we propose –where neuronal cilia deploy synaptic machinery to sense neurotransmitters– positions cilia as dynamic hubs for locally integrating neural circuit activity.

The discrete clusters of GluN1 on primary cilia raise important questions about signaling downstream of NMDAR in cilia. It is particularly interesting to consider the unique biophysical properties of NMDAR, which begins to conduct Ca^2+^ once the membrane potential rises above -40 mV^64^. In dendritic spines, NMDARs act as coincidence detectors for repeated synaptic firing and glutamate release, forming the molecular basis of long-term potentiation via CaMKII- and PKC-mediated remodeling of the postsynaptic compartment^65^. In contrast to dendritic spines, where the resting membrane potential is typically -65 mV, the ciliary membrane potential is reported to be 30 mV higher than that of the plasma membrane^66^. NMDARs on the ciliary membrane of neurons may thus be uniquely primed for activation by glutamate, making the hypothesis of a hypersensitive ciliary NMDARs detecting signals from local glutamatergic synapses particularly compelling. Similarly, astrocytes express NMDARs with reduced sensitivity to Mg^2+^ to sense synaptic glutamate efflux^67,68^.

Remarkably, GluN2B was recently detected on cilia of cultured neurons^69^, suggesting that GluN1 may assemble functional NMDA receptor heterotetramers with GluN2B on cilia. Once ciliary NMDAR permits Ca^2+^ entry, the presence of PKCδ and CaMKV in our brain cilia proteome (**Table S2**) suggest that these kinases may remodel the ciliary proteome. Beyond Ca^2+^ binding to effector molecules inside cilia, the flux of Ca^2+^ through NMDAR may further depolarize the ciliary membrane^70^. The presence of the voltage-gated Na^+^ channel Na_V_1.1 (**Fig. 5O**), and the Na^+^/K^+^ pump (**Fig. 5B**) in neuronal cilia reinforces the hypothesis of the cilium as a privileged electrical compartment. The best-characterized mechanism of cilium-based signaling in neurons comes from photoreceptors and olfactory receptor neurons^71^, where electrical signals initiated at the ciliary membrane propagate to the rest of the cell. Could primary cilia in other neurons also relay signals to the cell via changes in membrane potential? In line with this idea, activation of the ciliary calcium channel PKD2L1 was shown to increase the frequency of action potentials in hippocampal neurons^72^. The electrical output of neuronal cilia may thus tune the excitability of cortical neurons, positioning cilia as integrators of local synaptic signaling.

### Ciliary Adhesion Molecules and Neurite Regulation

Our proteomic analysis uncovered a diverse array of adhesion molecules, including NCAM1 (**Fig. 6K**), synCAM2 (**Fig. 5Q**), and protocadherin 9 (**Fig. 6G**), in neuronal cilia. These molecules likely mediate interactions between cilia and neighboring structures, including synapses and axons. Notably, NCAM1 appears to dynamically cycle in and out of cilia, accumulating along cilia when retrieval was blocked in *Bbs4^-/-^* neurons (**Fig. 6K-M**). This suggests that neuronal cilia may establish transient adhesive contacts with synapses or other cellular structures, allowing molecular clusters to be locally retained while other molecules are cleared. Such cilia adhesions may influence neuronal connectivity and plasticity.

In addition to conventional adhesion molecules, we uncovered several metabotropic GPCR, namely GABA_B1_ (**Fig. 5N**), mGluR5 (**Fig. 5P**) and GPR158 (**Video S2**), all of which are class C GPCRs enriched at the postsynaptic compartment. A defining feature of class C GPCRs is their large extracellular domains, which primarily function as orthosteric ligand-binding sites. In recent years, these extracellular domains have been shown to form transcellular complexes with adhesion molecules. For instance, GPR158 forms a transsynaptic complex with the heparan sulfate proteoglycan glypican-4 (GPC4)^73^, a presynaptic organizer; and productive GPR158-GPC4 complexes promote presynaptic compartment assembly. These findings suggest that ciliary metabotropic GPCRs may participate in axo-ciliary synapse formation.

Beyond adhesion, several regulators of neurite outgrowth, such as neogenin, a receptor that mediates repulsive guidance cues via activation of RhoA and actin cytoskeleton remodeling^74–76^, were enriched in neuronal cilia (**Figs. S8F-J**). Their enrichment in retrieval-deficient cilia (**Figs. 6J and S8F-J**) suggests that these molecules continuously patrol the neuronal cilium, dynamically sensing and regulating contacts between cilia and axons.

### Mechanisms Governing Nanodomain Formation on Neuronal Cilia

The precise spatial organization of signaling and adhesion molecules within neuronal ciliary nanodomains raises intriguing questions about the underlying mechanisms. Our findings suggest that active retrieval mechanisms play a pivotal role, with the BBSome restricting the excessive accumulation of signaling and adhesion molecules and ensuring that nanodomains remain functionally distinct. In addition, scaffolding proteins, including Ankyrin-B^77^ and SHANK3^63^, may anchor signaling complexes to specific sites on the ciliary membrane, creating stable nanodomains. Septins, which compartmentalize membranes in other systems, may act as physical barriers within cilia, segregating nanodomains and preventing diffusion^78^. Finally, lipid microdomains enriched in cholesterol or phosphoinositides^1,79^ may contribute to ciliary nanodomain organization, as seen in other specialized cellular compartments.

### Neuronal Cilia in Neuropsychiatric Disorders

Our findings have significant implications for understanding the molecular basis of neuropsychiatric disorders. Several proteins identified in the brain cilia proteome, including GluN1 and GABA_B1_, protocadherin-9 and SynCAM2, and LRRC7, are associated with schizophrenia^80–82^, autism spectrum disorders^83–85^, and intellectual disability^86^. Querying the disease association database DisGeNET^87^ revealed strong associations between the brain cilia proteome and these disorders (**Table S7**). These disorders are characterized by disruptions in synaptic signaling and neural circuit connectivity, suggesting that cilia-mediated signaling may play a previously unrecognized role in their pathogenesis. Supporting with this idea, ciliary defects in *Bbs2^-/-^* mice lead to deficits in social dominance^88^ and deletion of the serotonin receptor 6, a ciliary GPCR, leads to anxiety and cognitive impairments^89^. Targeting cilia-based signaling pathways may thus represent a promising therapeutic avenue for neuropsychiatric disorders.

## Conclusions

Our study highlights the diversity and specialization of neuronal cilia, revealing their unique molecular composition and close association with synaptic structures. By linking ciliary proteins to neuropsychiatric disorders, we establish a molecular framework for understanding how cilia-mediated signaling contributes to brain function. These findings pave the way for future studies aimed at unraveling the molecular mechanisms of cilia-synapse interactions and harnessing cilia biology for therapeutic innovation.

### Limitations of the study

While our study establishes a foundational resource for neuronal cilia biology, it also raises several unanswered questions. First, the functional roles of synaptic receptors and adhesion molecules in cilia remain speculative. Electrophysiological recordings or calcium imaging in cilia could provide direct evidence for their involvement in signal transduction. Second, the spatial organization of ciliary proteins suggests the existence of nanodomains, but the mechanisms governing their formation and maintenance are largely unknown. Investigating the role of the BBSome and scaffolding proteins in establishing these domains will be critical. Finally, our proteomics study was focused on the cerebral cortex, and specifically pyramidal neurons; the specialization and diversity of the neuronal cilia proteome will need to be studied in other brain regions.

## ACKNOWLEDGMENTS

We thank Drs. Rick Brown, Markus H. Schwab and Val Sheffield for the *Ubc-cre/ERT2*, *Nex-cre* and *Bbs4* mice; Philip Beachy for the Shh-N-producing HEK293T cells; Dr. Junli Zhang for pronuclear injection; Drs. Tamara Caspary and Karolina Nitsche for helpful discussions of mouse transgenesis; Adriana Hurtado for mouse colony management; Yien-Ming Kuo for help with microscopy; Suling Wang for help with graphic design; and Drs. Arturo Alvarez-Buylla, Grae David, Jeremy Reiter, Christian Vaisse, Markus Delling, Nicolas Berbari, Nathalie Spassky, and Won-Jing Wang and all members of the Nachury lab for stimulating discussions. This work was funded by NIH (GM089933 and EY031462 to MVN), ADA (1-20-VSN-03 to MVN), the National Science and Technology Council, Taiwan (MOST-110-2917-I-564-010 to CHC), the Sandler Program for Breakthrough Biomedical Research (to CHC and MVN), the UCSF Valhalla Fellows Program (to MEP), and CPRIT (RR220032 to MK who is a CPRIT scholar in cancer research). This work was made possible, in part, by EY002162 - Core Grant for Vision Research and by the Research to Prevent Blindness Unrestricted Grant (MVN).

The mass spectrometry proteomics data have been deposited to the ProteomeXchange Consortium via the PRIDE partner repository with the dataset identifiers PXD067626 and PXD067892.

## AUTHOR CONTRIBUTIONS

Conceptualization, CHC and MVN; Methodology, CHC, VNT, CKM, MEP, MK, and MVN; Formal analysis, CHC, VNT, NRL, MEP, MK, and MVN; Investigation, CHC, NRL, and CKM; Writing – original draft, CHC and MVN; Writing – review & editing, CHC, VNT, NRL, CKM, MEP, MK and MVN; Visualization, CHC and MVN; Funding acquisition, CHC, MVN, MEP and MK; Supervision, CHC, MK, MEP and MVN.

---

**Figure S1.**
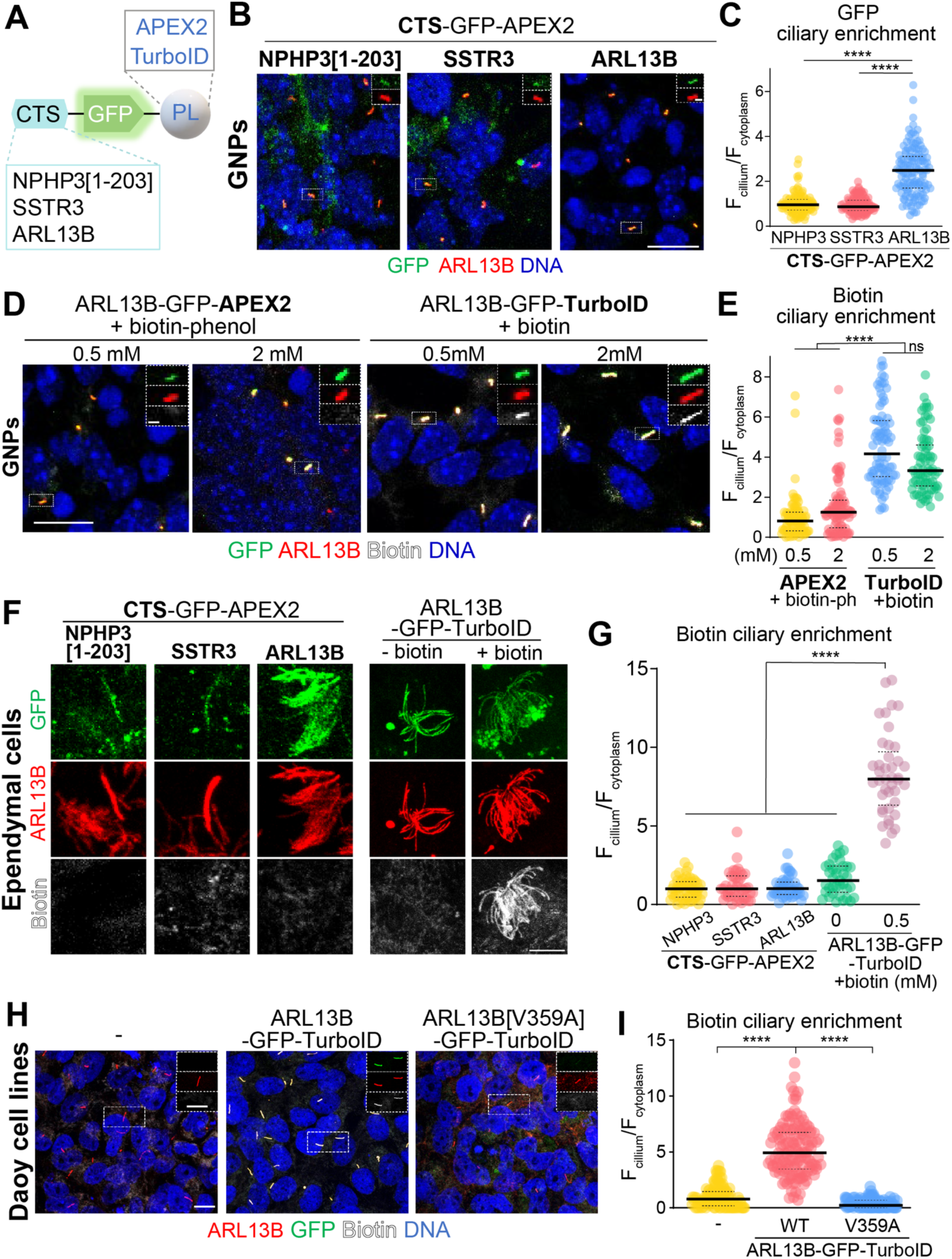
Engineering cilia-targeted proximity labeling tools in CNS cells. Related to Figure 1. **A.** Diagram of constructs tested for ciliary biotinylation. Ciliary-targeting sequences (CTS) were human NPHP3[1-203], mouse SSTR3, or human ARL13B. Each CTS was fused to GFP and either APEX2 or TurboID as the proximity labeling (PL) enzyme. **B.** Testing of CTSs in granule neuron progenitors (GNPs). GNPs from postnatal day 6 (P6) cerebella were transduced with lentiviruses expressing NPHP3[1-203]-GFP-APEX2 (left), SSTR3-GFP-APEX2 (middle), or ARL13B-GFP-APEX2 (right) and cultured for 4 days *in vitro* (DIV4). Cells were stained for GFP (green), ARL13B (red), and DNA (blue). Boxed regions with individual channels are displayed at the upper right of this panel with the following order (top to bottom): GFP, and ARL13B. Scale bars: 10 μm (main) and 1 μm (insets). **C.** Quantitation of ciliary enrichment of GFP fusions in GNPs. Scatter plots compare the ratio of GFP fluorescence in the cilium versus cytoplasm in each condition, normalized to the mean of the NPHP3[1-203]-GFP-APEX2 condition. The thick and dotted lines represent the median and interquartile range. *N* = 112-127 cilia per condition. **D.** Comparison of proximity labeling enzymes. GNPs were infected with lentiviruses carrying ARL13B-GFP-APEX2 or ARL13B-GFP-TurboID and treated at DIV4 for 1h with 0.5- or 2-mM biotin-phenol (for APEX2) or biotin (for TurboID). Cells expressing APEX2 were additionally treated with 1 mM H_2_O_2_ for 1 min prior to fixation. Cells were stained for GFP (green), ARL13B (red), biotin (white), and DNA (blue). Boxed regions with individual channels are displayed at the upper right of this panel with the following order (top to bottom): GFP, ARL13B, and biotin. Scale bars: 10 μm (main) and 1 μm (insets). **E.** Quantitation of ciliary biotin enrichment in GNPs. Scatter plots compare the ratio of biotin in the cilium versus cytoplasm under each condition, normalized to the mean of ARL13B-GFP-APEX2 cells treated with 0.5 mM biotin-phenol. The thick and dotted lines indicate the median and interquartile range. *N* = 75-98 cilia analyzed per condition. **F.** Testing proximity labeling fusions in primary ependymal cell cultures. Ependymal cells from P0 cerebral cortices were transduced at DIV6 with lentiviruses encoding the indicated constructs. At DIV21, APEX2-expressing cells were treated for 1h with 0.5 mM biotin-phenol and 1 mM H_2_O_2_ was added for 1 min prior to fixation, while TurboID-expressing cells were treated for 1h with biotin or left untreated before fixation. Cells were stained for GFP (green), ARL13B (red), and biotin (white). Scale bar: 5 μm. **G.** Quantitation of biotin enrichment in cilia of ependymal cells. Scatter plots compare the ratio of biotin fluorescence in the cilium versus cytoplasm in each condition, normalized to the mean of NPHP3-GFP-APEX2 cells treated with 0.5 mM biotin-phenol. The thick and dotted lines indicate the median and interquartile range. *N* = 37-52 per condition. **H.** Proximity labeling in Daoy medulloblastoma cells. Daoy cells (untransfected, left), or stably expressing either ARL13B-GFP-TurboID (middle) or ARL13B[V359A]-GFP-TurboID (right) were stained for ARL13B (red), GFP (green), biotin (white), and DNA (blue). Boxed regions with individual channels are displayed at the upper right of this panel with the following order (top to bottom): GFP, ARL13B, and biotin. Scale bars: 10 μm (main) and 5 μm (insets). **I.** Quantitation of ciliary biotin enrichment in Daoy cells. Scatter plots show the ratio of biotin fluorescence in the cilium versus cytoplasm, normalized to parental cells. Thick and dotted lines indicate the median and interquartile range. *N* = 99-108 cilia per condition. **C, E, G, I.** Statistical significance was determined by one-way ANOVA with post-hoc Dunn’s test (****, *p* < 0.0001; ns: not significant).

**Figure S2.**
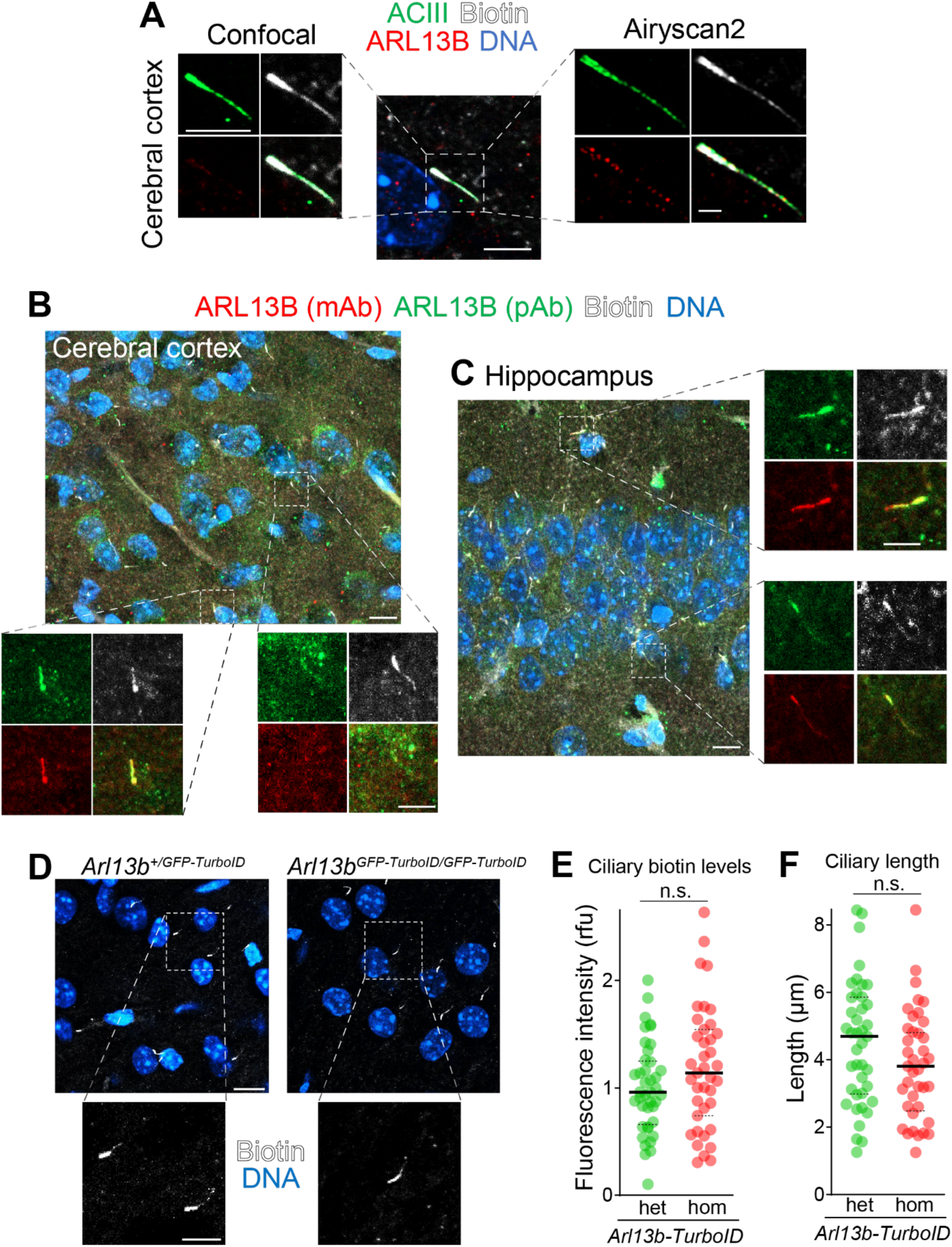
Low ciliary levels of ARL13B-GFP-TurboID in neurons saturate ciliary biotinylation. Related to Figure 2. **A.** Detection of biotin and ARL13B in pyramidal neuron cilia. Sections from the cerebral cortex of 3-month-old *Arl13b^+/GFP-TurboID^* mice injected with biotin were stained for ciliary markers (ACIII, green; ARL13B, red), biotin (white), and DNA (blue). A representative pyramidal neuron cilium (boxed) was imaged by conventional confocal microscopy (left panels) and by the Airyscan2 super-resolution module (right panels). The individual channels are presented in the following order (left to right and top to bottom): ACIII, Biotin, ARL13B and merge. Scale bars: 5 μm (main/confocal), 1 μm (Airyscan2). **B, C**. Detection of ARL13B in pyramidal neuron cilia. Cortical (**B**) and hippocampal Cornu Ammonis 1 (CA1) (**C**) sections of 3-month-old *Arl13b^+/GFP-TurboID^* mice injected with biotin were stained for ARL13B using monoclonal (mAb, red) and polyclonal (pAb, green) antibodies, and counterstained for biotin (white) and DNA (blue). Boxed regions are enlarged, and the individual channels are displayed in the following order (left to right and top to bottom): ARL13B (pAb), biotin, ARL13B (mAb), and merge. Scale bars: 10 μm (main), 5 μm (insets). **D-F.** Ciliary biotinylation in mouse cortices from heterozygous *Arl13b^+/GFP-TurboID^* (het) or homozygous *Arl13b^GFP- TurboID/GFP-TurboID^* (hom) mice. **D**. After 3 days of biotin administration in P21 mice, cerebral cortices were stained for biotin (white) and DNA (blue). The biotin signal in the boxed regions is shown at higher magnification in the bottom panels. Scale bars: 10 μm (main), 5 μm (insets). **E**. Scatter plots depicting the fluorescence intensity of the ciliary biotin signal relative to het cilia (RFU: relative fluorescence unit). **F.** Scatter plots of ciliary lengths measured from the biotin signals. Thick lines represent medians, and dotted lines indicate interquartile ranges. *N* = 39-41 cilia analyzed from 2 brains. Statistical significance determined by Mann-Whitney U test (ns: not significant).

**Figure S3.**
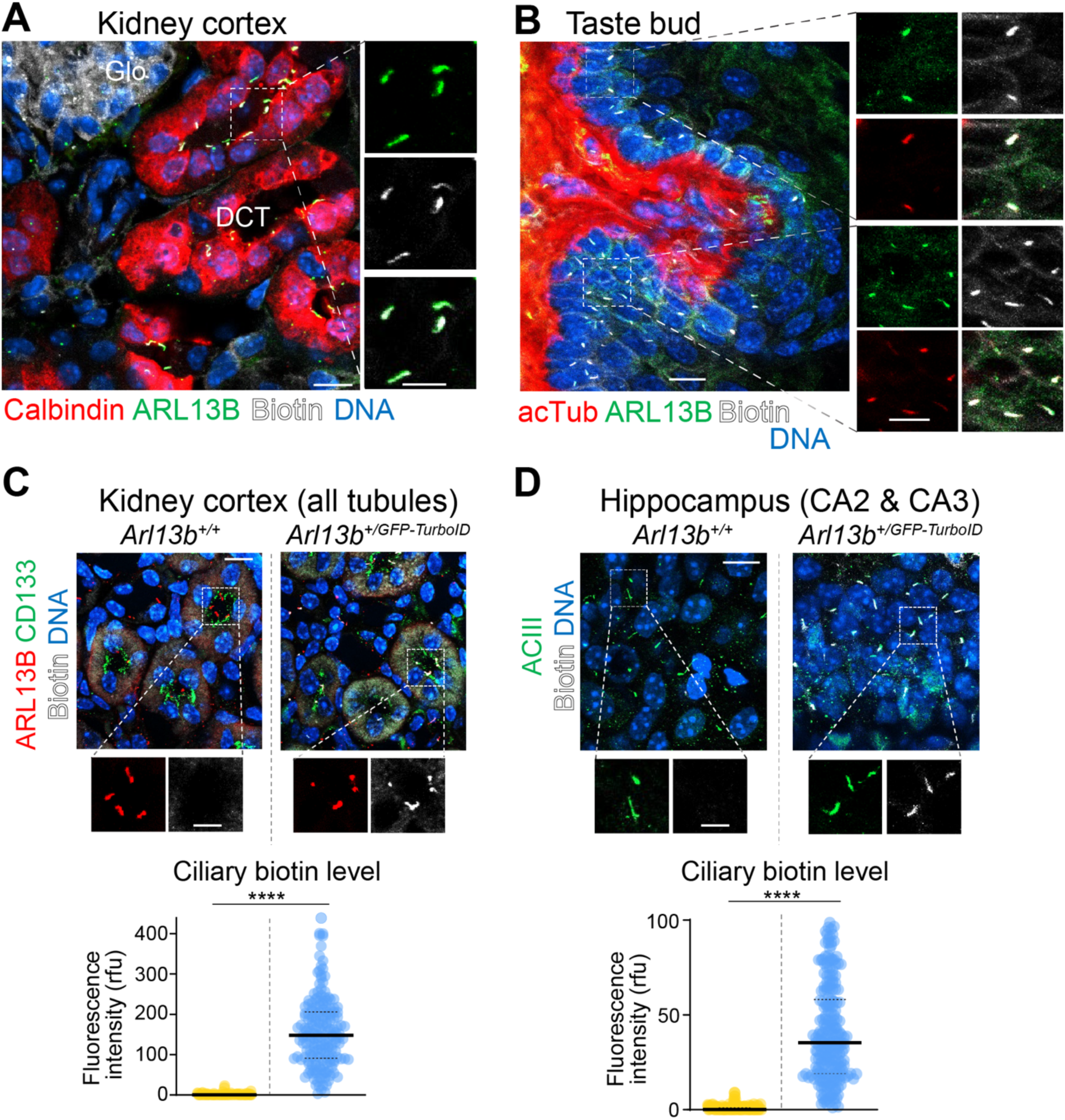
Cilia-specific biotinylation across tissues and biochemical purification of biotinylated proteins. Related to Figure 2. **A**. Biotin labeling in cilia of kidney tubular cells from *Arl13b^+/GFP-TurboID^* mice following biotin administration. Sections from the kidney cortex of P30 mice were stained for Calbindin D28K (red), ARL13B (green), biotin (white), and DNA (blue). The boxed region is shown at higher magnification on the right; channels are displayed in the following order: ARL13B, biotin, and merged. DCT: distal convoluted tubules; Glo: Glomerulus. Scale bars: 10 μm (main), 5 μm (insets). **B**. Cilia-specific biotinylation in the taste bud of *Arl13b^+/GFP-TurboID^* mice following biotin injection. Sections from the taste buds of 6-month-old mice were stained for acetylated tubulin (acTub, red), ARL13B (green), biotin (white), and DNA (blue). The boxed region is shown at higher magnification on the right; channels are displayed in the following order (left to right and top to bottom): ARL13B, biotin, acTub, and merge. Scale bars: 10 μm (main), 5 μm (insets). **C**. Cilia-specific biotinylation in the kidney of *Arl13b^+/GFP-TurboID^* mice. Kidney sections from 1-month-old wild-type (*Arl13b^+/+^*, left) or *Arl13b^+/GFP-TurboID^* (right) mice after 3 days of biotin administration were stained for a ciliary marker (ARL13B, red), a microvilli marker for delineating the apical lumen of kidney tubules (CD133, green), biotin (white), and DNA (blue). Boxed regions are enlarged below, displaying separated ARL13B (left) and biotin (right) channels. Scale bars: 10 μm (main), 5 μm (insets). Scatter plots depict the ciliary biotin fluorescence intensity normalized to the mean of *Arl13b^+/+^* cilia (RFU: relative fluorescence unit). Ciliary biotin fluorescence intensity in all renal tubules was measured. Thick lines represent the median, and dotted lines indicate the interquartile range. *N* = 177-188 cilia from 3 independent tissues. Statistical significance was determined by Mann-Whitney U test (****, *p* < 0.001). **D**. Cilia-specific biotinylation in the hippocampus of *Arl13b^+/GFP-TurboID^* mice. Hippocampal sections from 1-month-old wild-type (*Arl13b^+/+^*, left) or *Arl13b^+/GFP-TurboID^* (right) mice after biotin administration were stained for a cilia marker (ACIII, green), biotin (white), and DNA (blue). Boxed regions are enlarged below, showing separated ACIII (left) and biotin (right) channels. Scale bars: 10 μm (main), 5 μm (insets). Scatter plots depict the ciliary biotin fluorescence intensity normalized to the mean of *Arl13b^+/+^* cilia. Thick lines represent the median, and dotted lines indicate the interquartile range. *N* = 146-159 cilia from 3 hippocampus in CA2 and CA3 regions. Statistical significance was determined by Mann-Whitney U test (****, *p* < 0.001).

**Figure S4.**
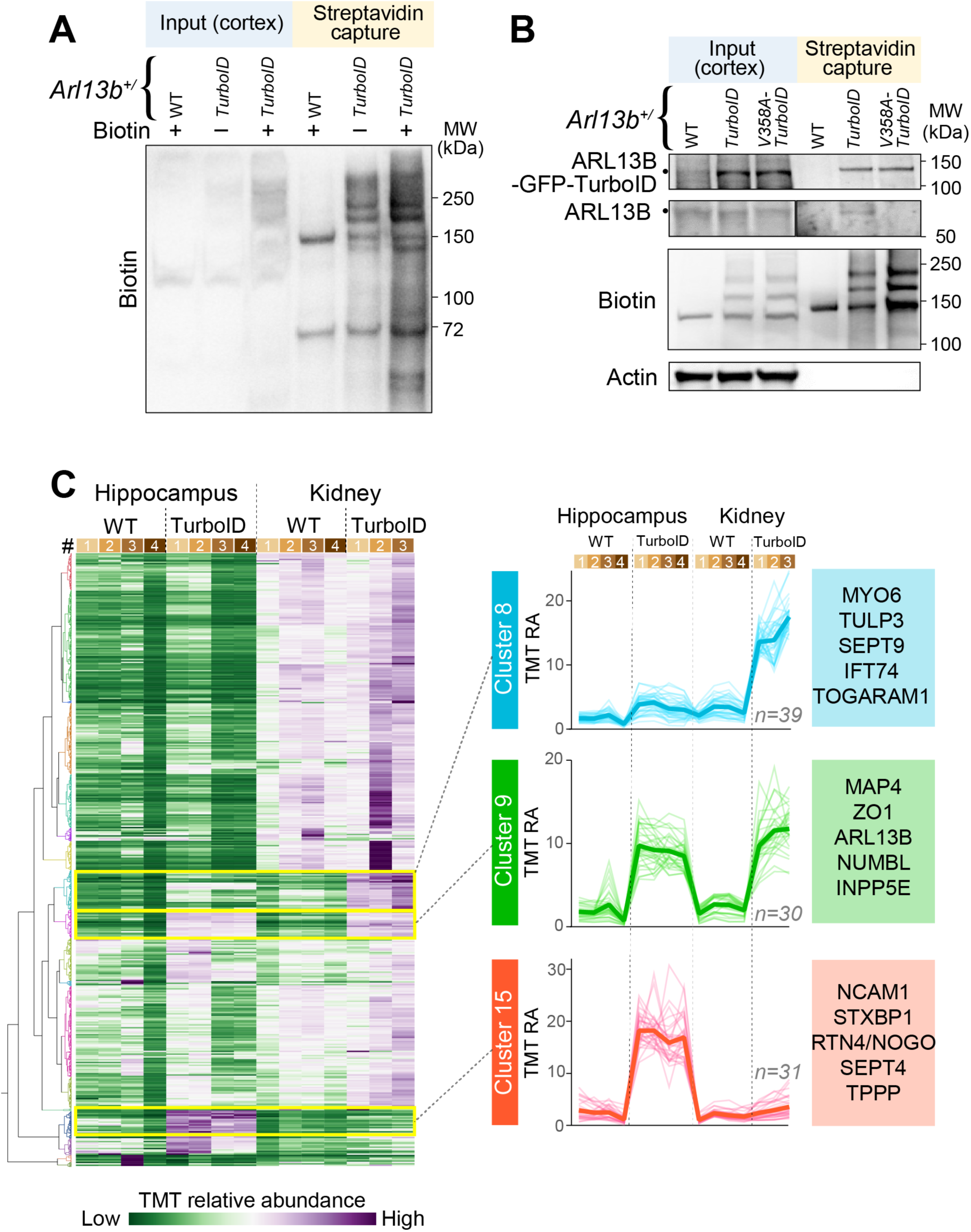
Comparative proteomics reveal distinct molecular signatures of kidney and hippocampal cilia. Related to Figure 2. **A**. Increased biotinylation upon biotin administration. Cerebral cortical lysates from P21 mice injected with either vehicle or biotin for 3 days were applied onto streptavidin beads. 5% inputs and 50% eluates were loaded for immunoblotting. The endogenous biotinylated proteins pyruvate carboxylase (130 kDa) and propionyl-CoA carboxylase ⍺ (70 kDa) are present in all samples. **B.** Biochemical validation of cilium-specific biotinylation. Cerebral cortices from 1-month-old WT (*Arl13b^+/+^*), TurboID (*Arl13b^+/GFP-TurboID^*), and V358-TurboID (*Arl13b^+/V358A-GFP-TurboID^*) mice were harvested after 3 days of biotin injection. Lysates were subjected to streptavidin capture and immunoblotted for ARL13B, β-actin and biotin. 1.67% inputs and 2.5% eluates were loaded for immunoblotting. The endogenous biotinylated protein pyruvate carboxylase is detected in all samples while the cytoplasmic protein, Actin, is not detectable in all eluates. Note that the ARL13B-TurboID and ARL13B[V358A]-TurboID protein can be detected in TurboID and V358-TurboID samples, indicating self-biotinylation^36^. **C**. Hierarchical clustering reveals distinct proteomic signatures of hippocampal and kidney cilia. After 7 days of biotin administration, biotinylated proteins were isolated from hippocampal and kidney tissues of wild-type (WT) and *Arl13b^+/GFP-TurboID^* (TurboID) mice and processed for Tandem Mass Tags (TMT) mass spectrometry analyses. Heatmap shows 1191 proteins across four WT hippocampus, four TurboID hippocampus, four WT kidney, and three TurboID kidney samples. Relative abundance (RA) was calculated by dividing TMT signal of each sample by the sum of TMT signals across all samples. The color scheme at the bottom indicates the RA scale. Line graphs correspond to the three boxed clusters. Thick lines represent the mean RA for each cluster, and thin lines denote individual proteins. Numbers indicate the total proteins in each cluster (see **Table S1** for dataset).

**Figure S5.**
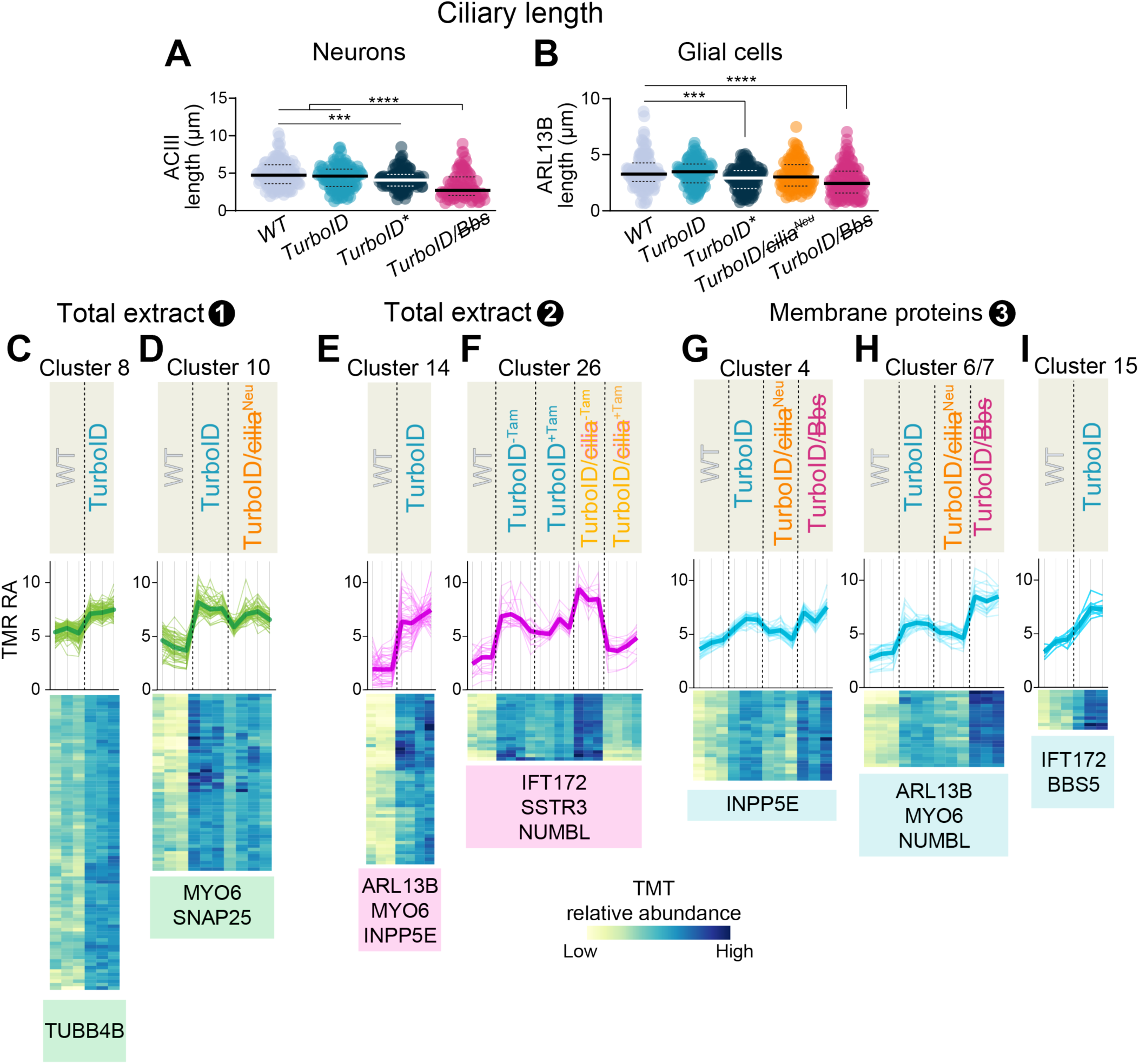
Ciliary lengths in control and experimental mice and analysis of ciliary proteomic clusters. Related to Figures 3-4. <u>Genotype abbreviations</u>: TurboID: *Arl13b^+/GFP-TurboID^* WT: *Arl13b^+/+^* TurboID†: *Arl13b*^+/*V358A-GFP-TurboID*^ TurboID/cilia: *Arl13b^+/GFP-TurboID^; Ubc-cre/ERT2; Ift88^fl/fl^* TurboID/cilia^Neu^: *Arl13b^+/GFP-TurboID^; Nex-cre; Ift88^fl/fl^* TurboID/Bbs: *Arl13b^+/GFP-TurboID^; Bbs4^-/-^* Tam: Tamoxifen **A, B**. Quantitation of ciliary lengths in neurons (**A**) and glial cells (**B**) across indicated genotypes. Scatter plots display ciliary length measured by ACIII (neurons) or ARL13B (glial cells). Thick lines represent the median, and dotted lines indicate the interquartile range. *N* = 105-227 cilia from 3 brains per genotype. Statistical significance was determined by one-way ANOVA with post-hoc Bonferroni test (***, *p* < 0.001; ****, *p* < 0.0001). See **Fig. 3A** for images. **C-I**. Clusters of ciliary proteins in three independent TMT-MS analyses. Total extracts (**C-F**) and membrane fractions (**G-I**) from dissected cortices of mice with indicated genotypes were purified on streptavidin beads and analyzed by TMT-MS as outlined in **Fig. 4E**. Line graphs depict relative abundance (RA), calculated via dividing TMT signal of each sample by the sum of the TMT signals across all samples. Thick lines show the average RA per cluster, and thin lines display individual proteins (the color scheme of each graph matches that of **Fig. 4B-D** with TMT experiment 1 in green, experiment 2 in magenta and experiment 3 in cyan). Heatmaps display the RA of each protein across indicated genotypes. The color scheme at the bottom indicates the RA scale. See **Table S2** for the detailed composition of each cluster.

**Figure S6.**
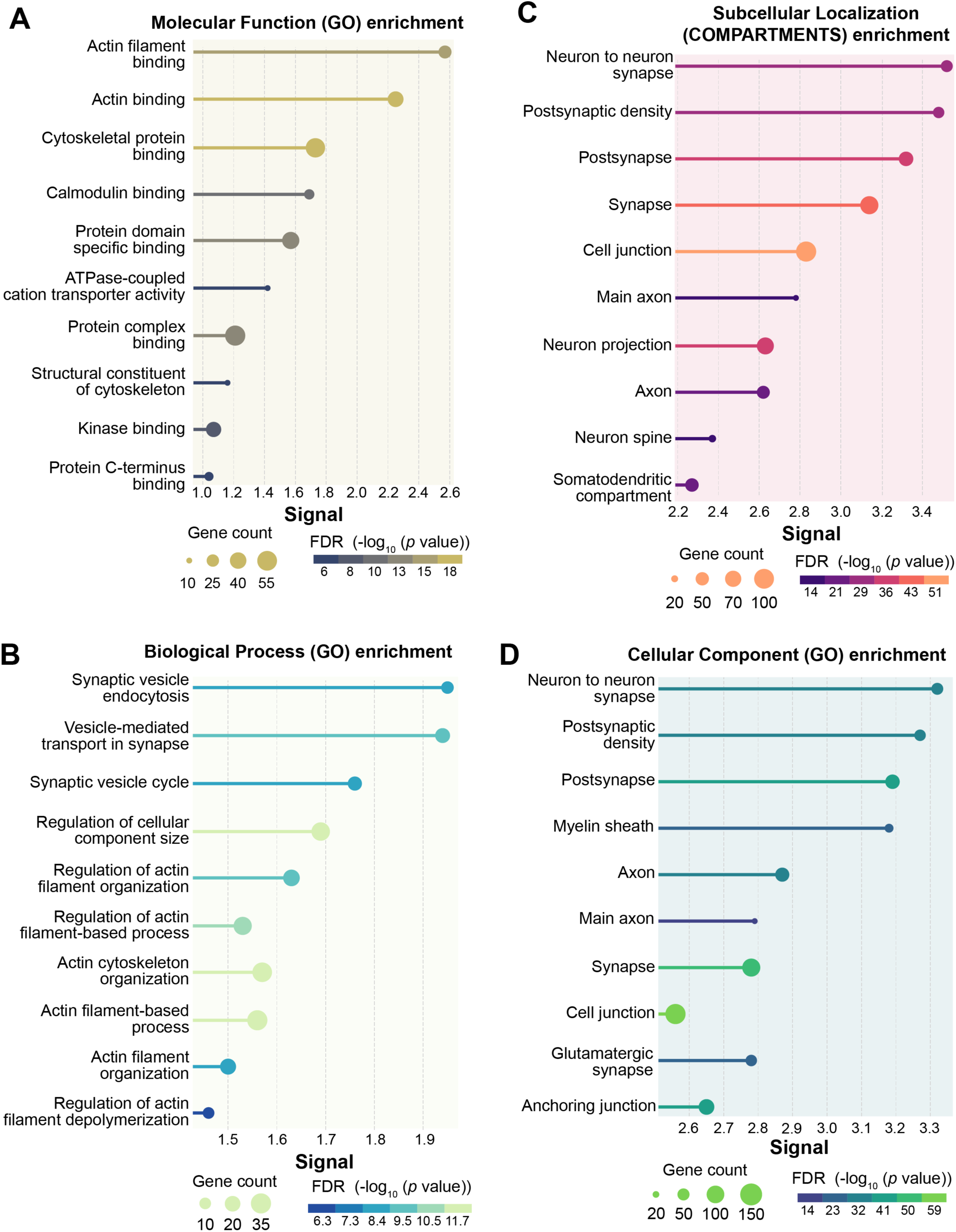
Gene ontology analysis of candidate brain cilia proteins. Related to Figure 5. **A-D**. The brain cilia proteome was subjected to a Gene Ontology (GO) enrichment analysis using g:Profiler^90^, STRING^91^, and SynGO^92^. Functional enrichment profiles are shown for molecular function **(A)**, biological process **(B)**, subcellular localization **(C)**, and cellular component **(D)**. False discovery rate (FDR) represents Benjamini–Hochberg-corrected *p*-values, indicating the significant of enrichment for each term. The signal is a weighted harmonic mean combining the observed/expected ratio and -log (FDR). Circle size corresponds to the number of genes associated with a given term, and the color scale represents the FDR. See **Table S3** for further analysis.

**Figure S7.**
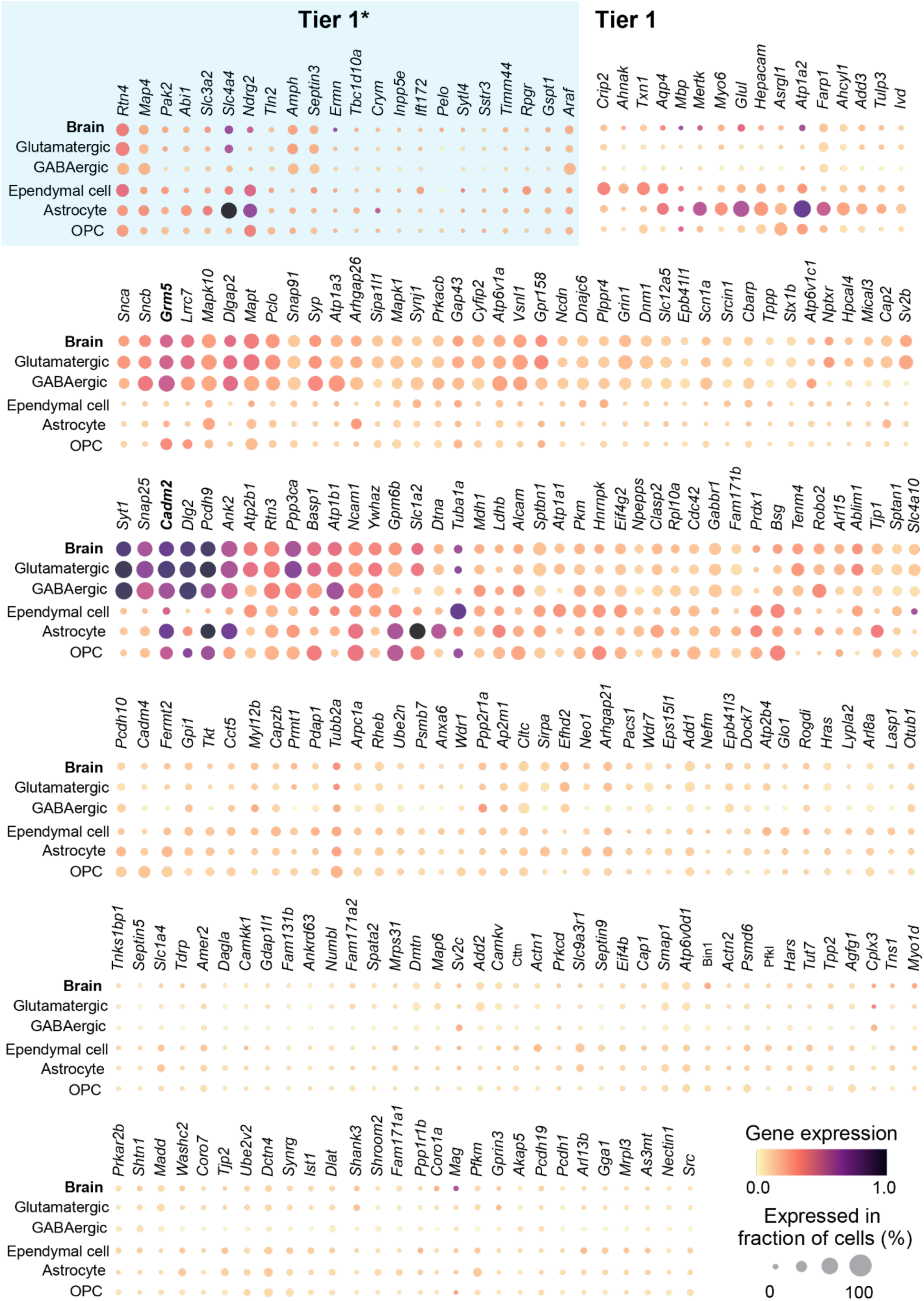
Candidate ciliary proteins sorted by gene expression in various ciliated CNS cell types. Related to Figure 5P-Q. Dot plots depict the expression of candidate ciliary proteins (separated by Tier1* [defined in **Fig. 6A, B**] and Tier1) across five ciliated cell types in the mouse brain. The data shown is extracted from the CZ CELLxGENE platform^55^, which integrates 16 single cell sequencing datasets comprising a total of 3.5 million cells. Each dot is color-coded by scaled average gene expression (ranging from 0 to 1) and sized according to the percentage of cells expressing the gene within each cell type. Genes are hierarchically ordered to highlight shared expression profile among the ciliated cell types. Note that proteins encoded by *Grm5* (mGluR5) and *Cadm2* (synCAM2) are validated in **Fig. 5P-Q**.

**Figure S8.**
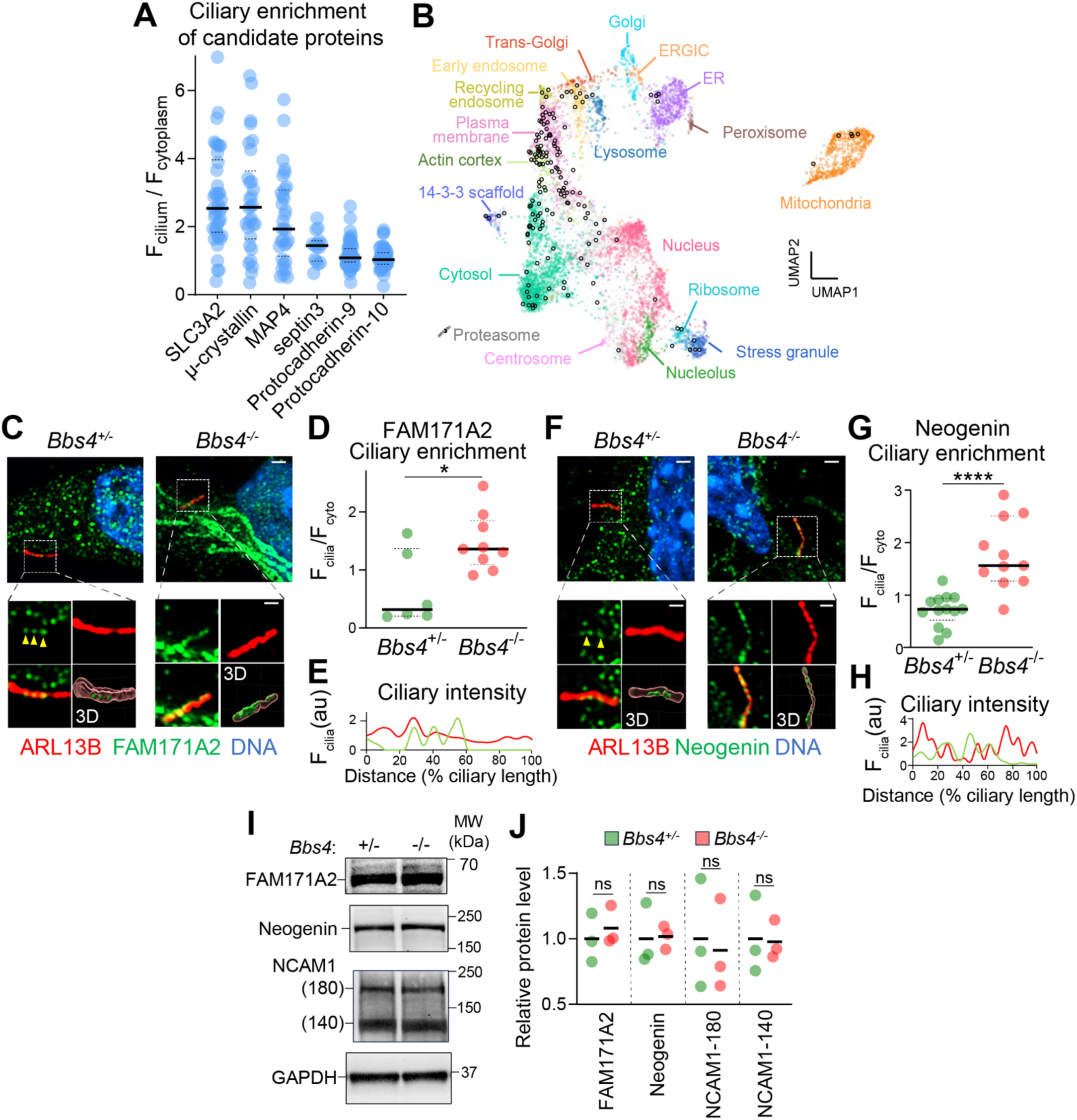
Analysis for cilia enrichment, organelle localization in Tier 1* and *Bbs4^-/-^*-enriched candidate proteins and the total protein levels in whole lysates from *Bbs4^+/-^* and *Bbs4^-/-^* brains. Related to Figure 6. **A** Quantitation of ciliary enrichment of candidate proteins in cultured neurons. Scatter plots show the ratio of candidate protein fluorescence in the cilium versus cytoplasm. Thick and dotted lines indicate the median and interquartile range. *N* = 15-68 cilia per condition. Related to **Fig. 6I**. **B**. Predicted subcellular localization of candidate cilia proteins. A two-dimensional Uniform Manifold Approximation and Projection (UMAP), derived from global organelle proteomic profiling^57^, preserves both local and global data structure and maps all Tier 1 candidate proteins to 20 distinct subcellular compartments. Each colored dot represents an individual protein while hollow circles indicate candidate cilia proteins. See **Table S5** for detailed analysis. **C, F**. Visualization of candidate protein localization to neuronal cilia. Cultured cortical neurons (DIV8) from E15.5 cerebral cortices were stained for candidate proteins (green), a ciliary marker (ARL13B, red), and DNA (blue). The boxed regions are enlarged to the bottom of each panel with the following order: candidate protein (top left), ARL13B (top right), merged channels (bottom left), and 3D reconstruction (Bottom right). Scale bars: 1 µm (main) and 500 nm (insets). **D, G**. Cilia enrichment of candidate proteins. Plots show the ratio of ciliary to cytoplasmic fluorescence for each candidate protein in individual cells. Thick and dotted lines represent the median and interquartile region. *N*=6-13 cilia scanned by Airyscan2 super-resolution module. Statistical significance was determined by the Mann-Whitney U test (*, *p* < 0.05; ****, *p* < 0.0001). **E, H.** Fluorescent intensity of candidate proteins along the shaft of cilium. Linescans reveal the fluorescence of candidate proteins (arbitrary unit, au) through the *Bbs4^+/-^* (green) and *Bbs4^-/-^* (red) cilia. The length of cilium is normalized to 100%. **I, J**. Candidate proteins enriched in *Bbs4^-/-^* cilia were immunoprobed in brain lysates. **(I)** Whole brain lysates from P30 *Bbs4^+/-^* and *Bbs4^-/-^* cerebral cortices were immunoblotted for FAM171A2, NEO1, NCAM1, and GAPDH. **(J)** Quantitation of protein levels in *Bbs4^-/-^* (red) relative to *Bbs4^+/-^* (green) cortices. Note that two isoforms of NCAM1 (140 and 180kDa) were quantified separately. Protein levels of *Bbs4^-/-^* samples are normalized to the mean of *Bbs4^+/-^* samples in each candidate protein. Thick lines mark the mean in each condition. Statistical significance was determined by unpaired *t* test (ns: not significant). *N* = 3 animals.

**Figure S9.**
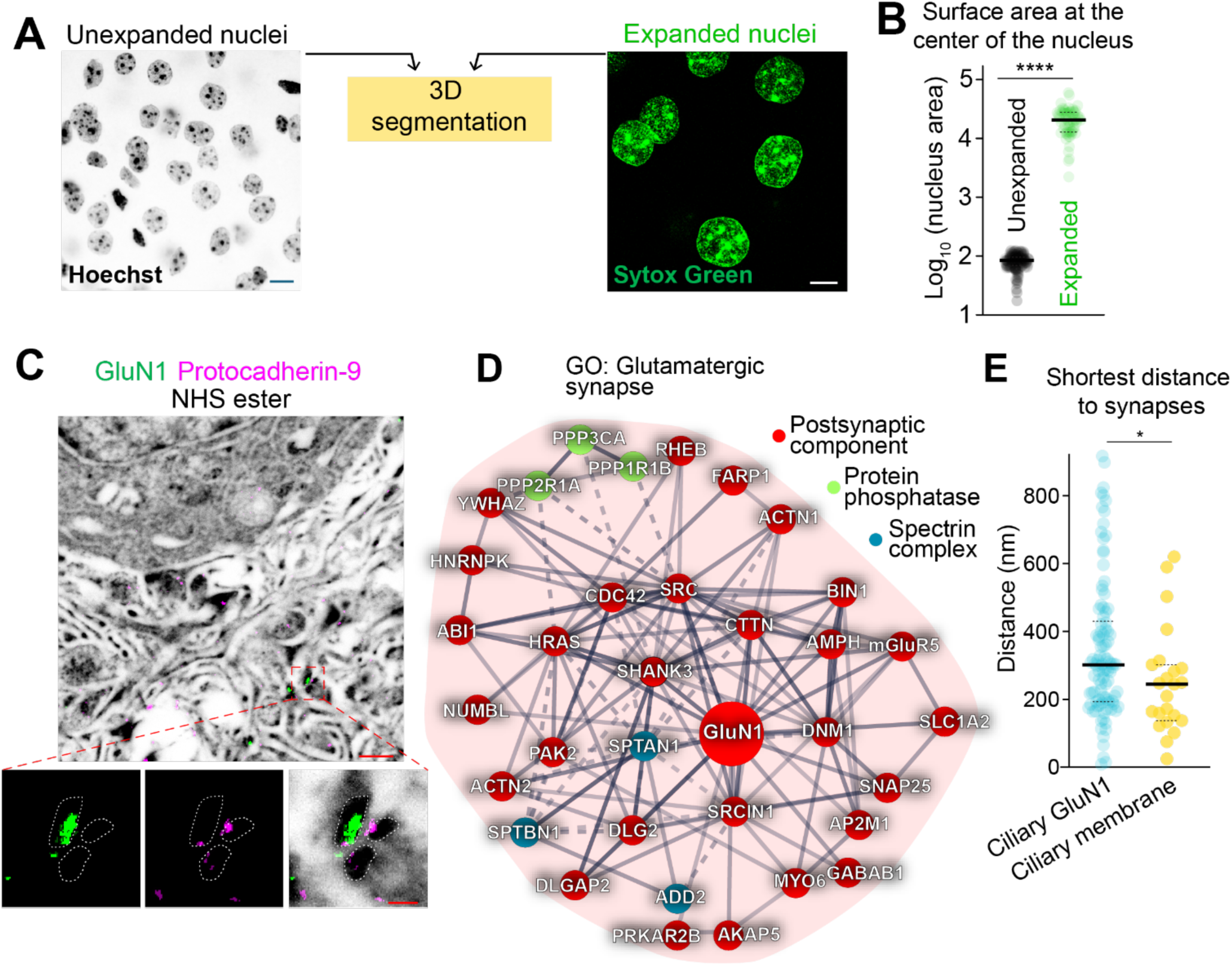
Pan-ExM-t reveals the ultrastructure of neuronal cilia and synapses. Related to Figure 7. **A.** Automated quantitation of the mid cross-sectional area of neuronal nuclei in expanded versus unexpanded cerebral cortices from 2-month-old wild-type (WT) mice. Samples subjected to expansion microscopy (ExM) were stained with Sytox Green, and unexpanded tissues were stained with Hoechst (grey), followed by automated 3D segmentation and counting^93^. Images shown here were processed with maximal intensity projection covering a given nucleus. Scale bars: 100 µm (expanded) and 10 µm (unexpanded). **B.** Scatter plot of the mid cross-sectional area of nuclei in unexpanded (*N* = 244) and expanded (*N* = 75) samples. The surface area of each nucleus in the largest cross-section is shown on a log scale, with thick lines indicating median and dotted lines showing interquartile ranges. Statistics: Mann-Whitney U test (****, *p* < 0.0001). The averaged cross-sectional areas are 82.11 µm^2^ (unexpanded) and 21628.34 µm^2^, corresponding to a linear expansion factor of 16.2. **C.** Identification of neuronal synapses in pan-ExM-t. P60 cerebral cortices were expanded and stained for the NMDA receptor (GluN1, green), and for the synaptic molecules PCDH9 (magenta), with a fluorophore-conjugated NHS ester (inverted greyscale). Boxed regions with separate channels are enlarged at the bottom of this panel with the following order (left to right): GluN1, PCDH9, and merged with NHS-ester staining. The envelope of the pre- and post-synaptic densities seen in the NHS ester channel are marked by dotted lines. Scale bars (adjusted by expansion factor): 500 nm (main) and 100 nm (insets). **D.** STRING analysis of interactions between candidate brain cilia proteins that match the GO term ‘glutamatergic synapse’. Protein names are colored for postsynaptic components (red), protein phosphatases (green), and spectrin complexes (blue). Lines thickness indicates the interaction confidence. **E.** Scatter plot depicting the shortest distance between 3D-rendered synaptic GluN1 spots to ciliary GluN1 molecules (cyan, *N* = 102) and to the nearest segmented cilia (yellow, *N* = 23). Thick lines indicate median and dotted lines show interquartile ranges. Statistics: Mann-Whitney U test (*, *p* < 0.05).

## STAR★METHODS

## KEY RESOURCES TABLE

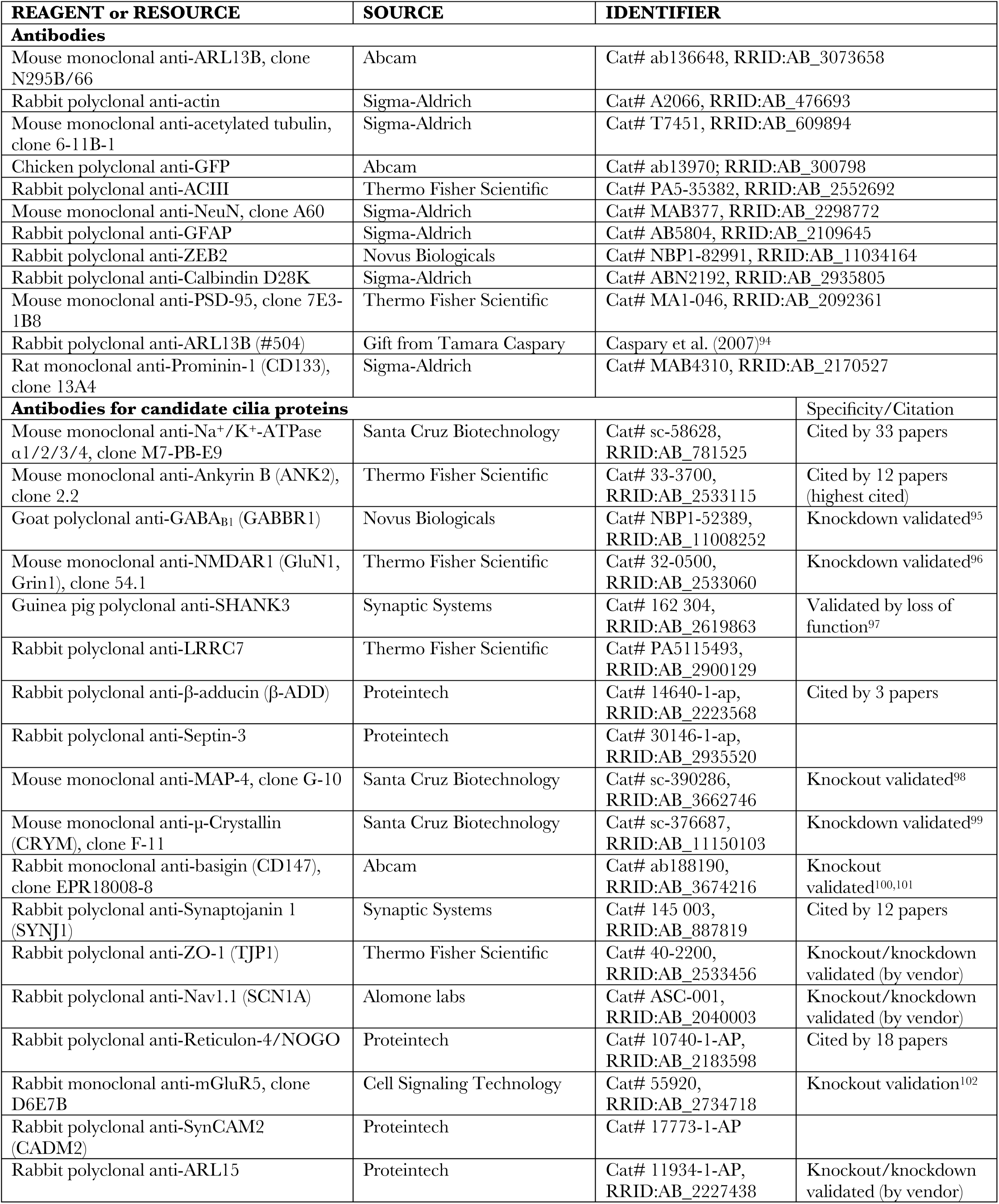
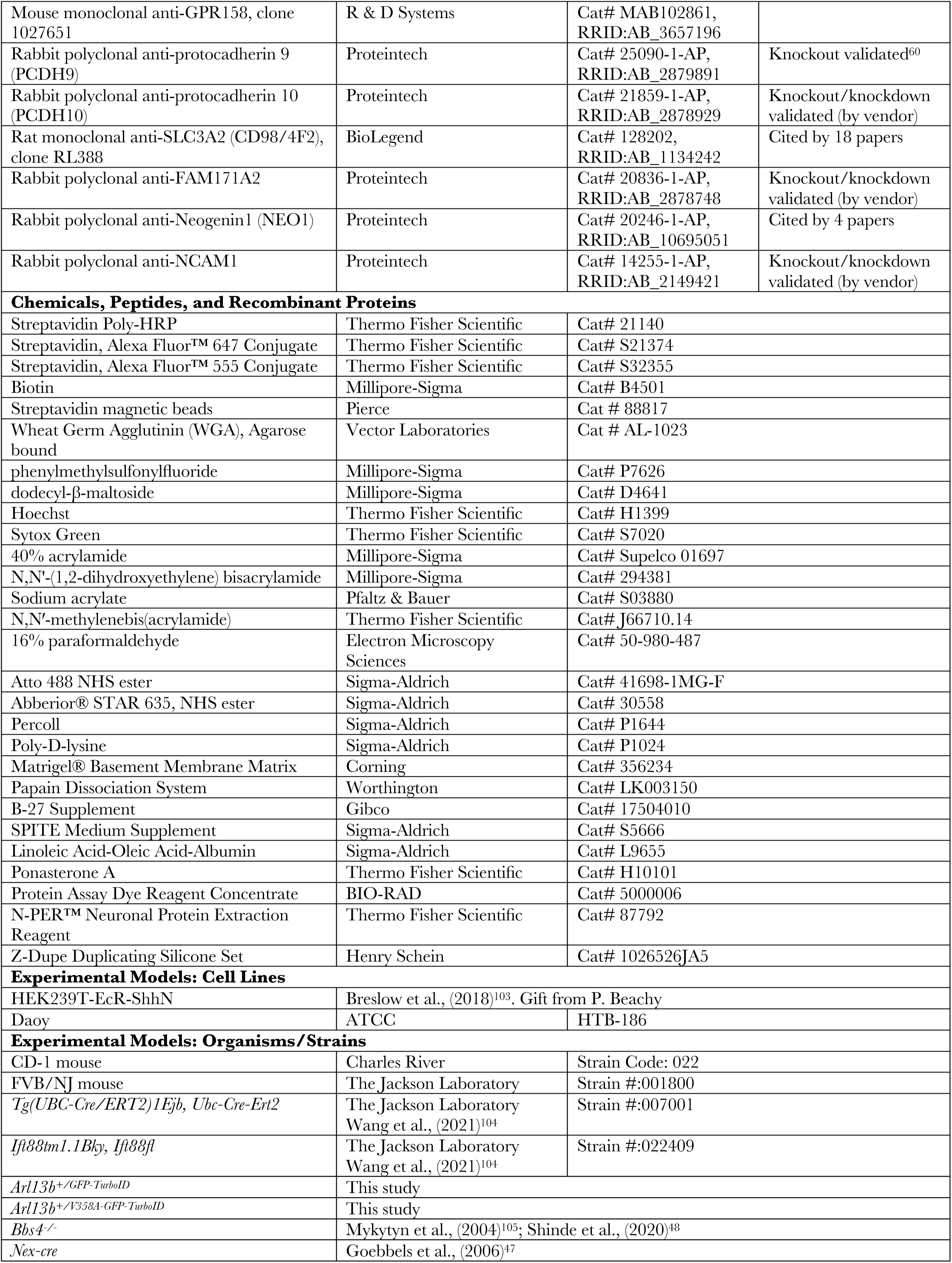
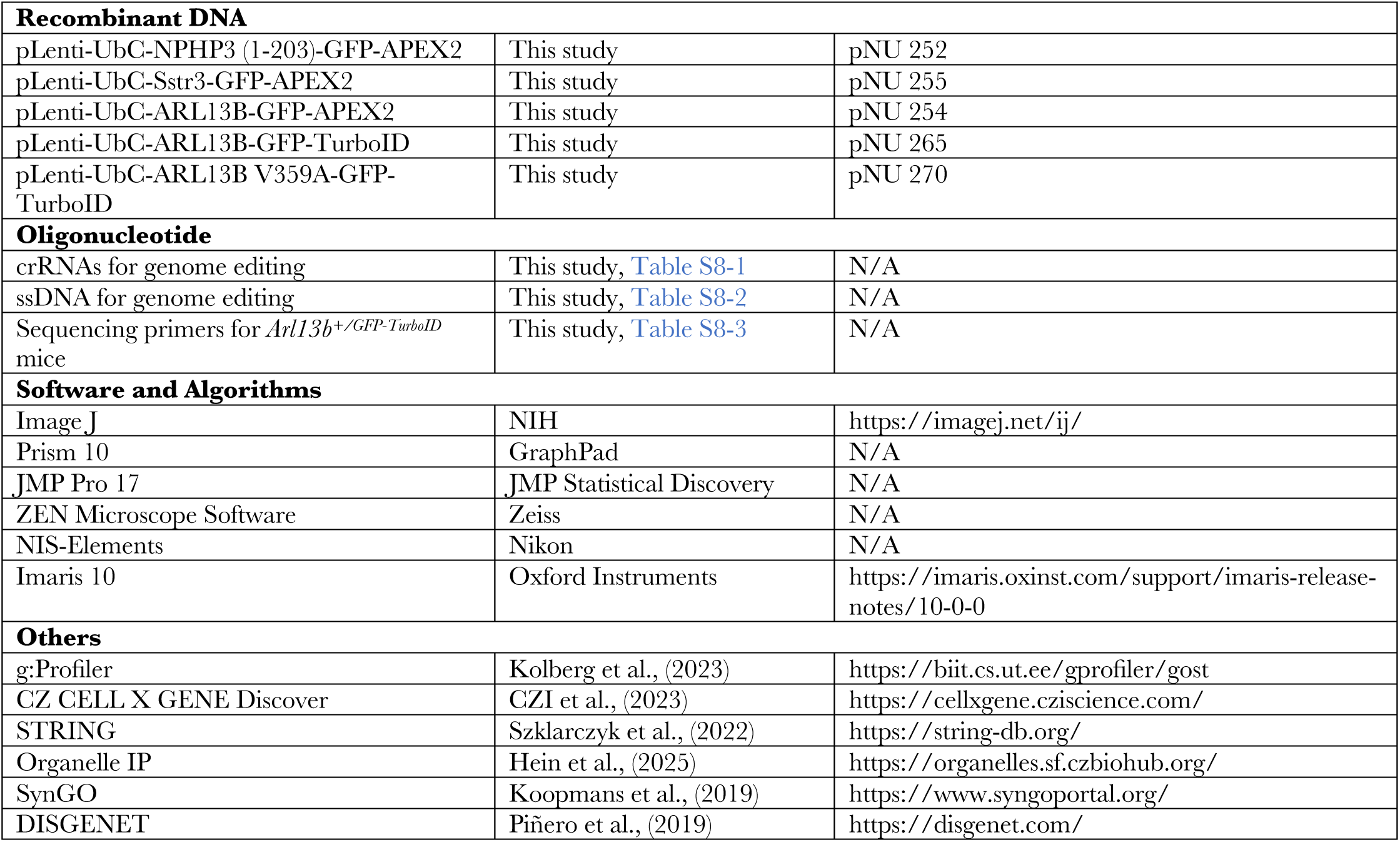

## CONTACT FOR REAGENT AND RESOURCE SHARING

### Lead contact

Further information and requests for resources and reagents should be directed to and will be fulfilled by the lead contact Maxence Nachury.

### Materials availability

This study has generated plasmids and cell lines, which are listed in the Key resources table. These reagents will be made available upon request.

### Data and code availability

The mass spectrometry proteomics data have been deposited to the ProteomeXchange Consortium (http://proteomecentral.proteomexchange.org) via the PRIDE^106^ partner repository with the dataset identifier PXD067626/PXD067892 and DOI: 10.6019/PXD067626. All the other data reported in this paper will be shared by the lead contact upon request. This paper does not report original code. Any additional information required to reanalyze the data reported in this study is available from the lead contact upon request.

## EXPERIMENTAL MODEL AND SUBJECT DETAILS

### Animal Husbandry

Primary cultures of GNPs, ependymal cells, and cortical neurons were conducted from CD-1 mice, which were purchased from Charles River. Transgenic mouse strains, including *Tg(UBC-Cre/ERT2)1Ejb* and *Ift88tm1.1Bky*, were obtained from the Jackson Laboratory^104^. The *Arl13b^+/GFP-TurboID^* founder mouse was initially generated in FVB/NJ and subsequently backcrossed to the CD-1 background for three generations. A maximum of five adult mice were housed per cage. Breeding cages contained one male and up to two females. Both male and female mice were used in all studies. To knockout *Ift88* postnatally, tamoxifen was injected subcutaneously into *Arl13b^+/GFP-TurboID^; Ubc-cre/ERT2; Ift88^fl/fl^* and *Arl13b^+/GFP-TurboID^; Ift88^fl/fl^* (control) pups at P5 for five consecutive days, followed by a three-week waiting period prior to conducting proteomics or immunostaining. All procedures adhered to the IACUC guidelines of UCSF. This project, ‘‘Molecular and Proteomic Studies of the Cilia,’’ was reviewed and approved by the IACUC at UCSF (AN195589).

### Cell Lines

Daoy cells (ATCC, HTB-186), derived from human desmoplastic cerebellar medulloblastoma, were cultured in Eagle’s Minimum Essential Medium (Corning #10-009-CV) supplemented with 10% fetal bovine serum (FBS, from Gemini Bio-products #100-106) and 100 units/mL of penicillin and streptomycin (P-S, from Gibco #15140-122). HEK293T-EcR-ShhN cells, provided by P. Beachy, were cultured in Dulbecco’s Modified Eagle’s Medium (DMEM, Gibco #11965092) supplemented with 10% FBS, 2 mM L-glutamine (Gibco #25030081), and antibiotics. To produce Shh-N conditioned media, HEK293T-EcR-ShhN cells were cultured in GNP culture media and stimulated with ponasterone A (5 µM) for 48h. The supernatant was collected and filtered by a 0.2 µm PVDF syringe filter (Whatman #9913-2502). A range of dilutions of Shh-N-conditioned media was tested on GNP cultures and the optimal dilution was chosen based on a GNP ciliation rate of 90%. All cell lines used in this study were tested monthly for mycoplasma contamination using the MycoAlert system (Lonza).

### Primary Culture

Primary cultures of purified GNPs from P6 cerebella were conducted as previously described^107^. Briefly, cerebella were digested using the papain dissociation system (Worthington) according to the manufacturer’s instructions. Dissociated cells were purified via a Percoll gradient, and glial cells were removed by pre-plating. Purified GNPs were cultured in Neurobasal medium (Gibco) supplemented with 100 units/mL P-S, 2 mM L-glutamine, 0.45% D-glucose, 2% B-27 (Gibco), 1% Linoleic Acid-Oleic Acid-Albumin solution (Sigma-Aldrich), 100 µM N-acetylcysteine, 1% SPITE medium (Sigma-Aldrich), and 2% Shh-N conditioned medium for 3 days. Lentivirus carrying genes of interest was added on the first day of primary culture.

Primary cultures of cortical neurons were conducted as previously described^108^. Briefly, cerebral cortices from E13.5-E15.5 embryos were dissociated using the papain dissociation system, and cell clumps were removed using cell strainers (Falcon #352350). Cells were cultured in Neurobasal medium supplemented with 100 units/mL P-S, 2 mM L-glutamine, 0.45% D-glucose, and 2% B-27 (Gibco) for 7-9 days.

Primary cultures of ependymal cells were conducted as previously described^109^. Briefly, dissociated cells from P0 cerebral cortices were cultured in 25 cm² flasks in DMEM-GlutaMAX medium (Gibco) supplemented with 10% FBS and antibiotics for 3-5 days. After reaching confluence, neurons and oligodendrocytes were removed by shaking the flasks at 250 rpm on an orbital shaker at room temperature overnight. Subsequently, 20 µl of ependymal cells (10^7^ cells/ml) were placed in the center of a 12mm circle coverslip (Fisher Scientific #12541001) for 1 day and cultured in media without FBS for an additional 2 weeks.

Coverslips for culturing GNPs and cortical neurons were pre-treated with poly-D-lysine (Sigma-Aldrich) and Matrigel (Corning) before seeding cells.

## METHOD DETAILS

### Plasmids

Sequences encoding human NPHP3[1-203] fused to GFP-APEX2, mouse SSTR3 fused to GFP-APEX2, human ARL13B fused to GFP-APEX2, and human ARL13B fused to GFP-TurboID were subcloned into the pLenti6-UbC-V5-DEST vector (Thermo Fisher Scientific, Cat. # V49910). Site-directed mutagenesis was performed to generate the pLenti-UbC-ARL13B[V359A]-GFP-TurboID plasmid using Pfu Turbo DNA polymerase according to manufacturer’s instructions (Agilent, Cat. # 200518).

### Virus Production and Transduction

Lentiviruses carrying NPHP3[1-203]-GFP-APEX2, SSTR3-GFP-APEX2, ARL13B-GFP-APEX2, ARL13B-GFP-TurboID, and ARL13B[V359A]-GFP-TurboID were produced as described in manufacturer’s protocol (PEG-it, System Biosciences). Briefly, lentiviral vectors along with the plasmid of interest were transfected into HEK293T cells using Lipofectamine 2000 (Thermo Fisher Scientific) according to the manufacturer’s protocol. Supernatants containing viral particles were concentrated using PEG-it Virus Precipitation Solution (System Biosciences) and re-suspended in PBS. Lentiviruses were added to dissociated cells on the first day of primary culture and washed away the following day.

### Proximity labeling experiments *in vitro*

APEX-based proximity labeling was performed as described^34^. Briefly, primary cells transduced with APEX2 fusions were cultured in their indicated medium supplemented with 0.5 mM (or specified concentration) biotin-phenol for 1 hr. Next, cells were incubated with 1 mM H_2_O_2_ in culture medium for 1 minute at room temperature, then washed three times in quenching buffer, consisting of 10 mM sodium ascorbate, 10 mM sodium azide, and 5 mM Trolox in PBS. For proximity labeling using TurboID, primary cells expressing TurboID were incubated in medium containing 0.5 mM (or specified concentration) biotin for 1 hr and washed with cold PBS before fixation.

### Immunostaining and Imaging

Primary cells or Daoy cells were fixed with 4% PFA in PBS for 15 minutes, followed by treatment with pre-chilled methanol for 5 minutes at -20°C. Cells were permeabilized in PBST (0.2% Triton X-100 in PBS) for 30 minutes and blocked with PBS containing 5% normal goat serum (NGS) and 5% bovine serum albumin (BSA) for 1h. Cells were incubated with primary antibodies, as listed in the Key Resources Table, and secondary antibodies (Alexa Fluor, Invitrogen or Jackson ImmunoResearch Labs) for 2 hr each at room temperature. Cells were counterstained with Hoechst (Thermo Fisher Scientific) and coverslips mounted onto slides using Fluoromount-G (Electron Microscopy Sciences). For Airyscan2 super-resolution imaging, cells were mounted in ProLong Antifade (Thermo Fisher Scientific).

For immunostaining of tissues, mice were transcardially perfused with 4% PFA in PBS, and tissues were post-fixed in 4% PFA overnight. Coronal sections (100 µm) were prepared using a Sliding Microtome (Thermo Fisher Scientific). We applied heat-mediated antigen retrieval with citrate buffer (10 mM Sodium citrate, 0.05% Tween 20, pH 6.0) in PBST-permeabilized samples, which were then incubated with primary antibodies (as listed in the Key Resources Table) for 2 days, followed by incubation with secondary antibodies for 2 hr. Tissues were counterstained, mounted as described above, and imaged on either an LSM900 microscope equipped with Airyscan2 (Zeiss) or an AX/AX R NSPARC system (Nikon). For imaging of tissues, a 40x/1.4NA oil-immersion objective (Zeiss) was used at 1024 x 1024-pixel resolution. For cultured cortical neurons, Airyscan2 super-resolution imaging was performed with a 63x/1.4NA oil-immersion objective, using a pixel size corresponding to 5-fold Nyquist sampling and 0.1µm optical sections.

### Image Analysis

For fluorescence intensity measurements in cilia of cultured neurons, sum-intensity projections were first created from consecutive Z-sections encompassing the entire cilium. A segmented line mask was then drawn along each cilium and transferred to the proteins of interest (POI) channel. The means of raw intensities in cilia (F _raw_) and adjacent cell-free areas (F _background_) were measured, and the final mean ciliary fluorescence intensities (F _cilia_) were calculated by the following formula: F _cilia_= F _raw_ – F _background_. For cilia enrichment analysis, the same procedure was followed, and an additional mask outlining the cell boundary was generated to measure the POI intensity in the cytoplasm (F _cytoplasm_). The cilia enrichment ratio was then calculated as (F _raw_ – F _background_) / (F _cytoplasm_ – F _background_).

For fluorescence intensity measurements in ARL13B-labeled cilia from tissues (**Figs. 2B, 2I, 2K, 3E, and S3C**), cilia were segmented in 3D using the Threshold method and FilterObjects functions in the GA3 platform (NIKON NIS software). The total biotin intensity (F _cilia_) and the number of voxels (V _cilia_) in each segmented cilium were recorded. To estimate the background intensity, each segmented cilium was dilated by a 3-pixel radius in three dimensions; the total biotin intensity (F _dilate_) and the voxel count (V _dilate_) of the dilated cilia were then measured. The averaged background intensity (F _background_) of each cilium was determined using the following formula: F _background_= (F _dilate_ – F _cilia_)/ (V _dilate_ – V _cilia_). Finally, the mean biotin intensity for each cilium was calculated by subtracting its background intensity from the mean intensity: F_total_/V_cilia_ – F_background_.

For fluorescence intensity measurements in ACIII-labeled cilia from tissues (**Figs. 2B, 2J, 3C, and S3D**), Denoise.ai (NIKON NIS-Elements) was applied to remove Poisson-distributed shot noise^110^, thus eliminating the granular background pattern while preserving ciliary ACIII signals. To verify the quality of segmentation, all the segmented images were aligned to the raw images. Measurements of biotin intensity within ACIII-labeled cilia were then carried out using the same approach as for ARL13B-labeled cilia. Quantitated data were plotted, and statistical tests were performed in GraphPad Prism, as indicated in the respective figure legends.

For linescan analysis, longitudinal fluorescence intensities of the POI were measured in ImageJ by a line extending along the ciliary length. Pixel intensities for each cilium were plotted against the percentage of total length, from 0% at the base to 100% at the tip.

### Generation of *Arl13b*^+/*GFP-TurboID*^ and *Arl13b*^+/*V358A*-*GFP-TurboID*^ Mice

Mouse genome engineering was conducted at the Transgenic Gene Targeting Core (TGTC) at the Gladstone Institute. We followed the Easi-CRISPR protocol^111^ to generate the *Arl1-GFP-TurboID* and *Arl13[V358A]-GFP-TurboID* alleles in mice. 4-week-old FVB/NJ female mice were injected with pregnant mare serum gonadotropin (PMSG) and human chorionic gonadotropin (HCG). Female mice with timed hormone induction were then bred with single-housed 10-week-old male mice. After dissection of the female reproductive system, fertilized oocytes and zygotes were collected and transferred to TGTC, where pronuclear injection of the ctRNP complex, containing crRNA (**Table S8**), tracrRNA (IDT), and Cas9 proteins (IDT), along with ssDNA (**Table S8**), was performed. Injected embryos were implanted into surrogate female mice and collected 17 pups at P21. Genomic DNA was extracted from each animal and several fragments spanning exon 10 of *Arl13b* to *GFP-TurboID* were amplified and subjected to Sanger sequencing. One out of 17 mice possessed a *GFP-TurboID* cassette in frame with the last amino acid of *Arl13b*. This animal was backcrossed to CD1 mice for three generations to remove possible extragenic mutations. Mice were genotyped with triplex primers (**Table S8**), where Arl13b^WT^ and Arl13b^GFP-TurboID^ alleles yielded products of 316 and 585 bp.

To generate the *Arl13b[V358A]* variant, we mated 4-week-old *Arl13b^GFP-TurboID/GFP-TurboID^* female mice with 10-week-old *Arl13b^GFP-TurboID/GFP-TurboID^* male mice and harvested fertilized oocytes and zygotes as described above. Zygotes were electroporated with the ctRNP complex and ssDNA (**Table S8**). Mice harboring the V358A mutation on the *Arl13b^GFP-TurboID^* allele were backcrossed to wild-type (*Arl13b^+/+^*) CD1 mice and the offsprings with the V358A mutation on the *Arl13b^GFP-TurboID^* allele were selected as *Arl13b^+/V358A-GFP-TurboID^* founder mice. This animal was backcrossed to CD1 mice for three generations to remove possible extragenic mutations. We observed no gross abnormalities in mice heterozygous for *Arl13b[V358A]-TurboID*, consistent with prior studies of mice possessing one *Arl13b[V358A]* allele^37^. Genotyping of the *Arl13b[V358A]* mutation was performed as described^37^.

### Biotin injection, streptavidin capture and immunoblotting

Mice were injected with 5 µL/g (body weight) biotin (24.4 mg dissolved in 1mL biotin, then mixed with 1mL PBS) intraperitoneally for 7 consecutive days starting from P30. The day after the last biotin administration, mice were transcardially perfused with PBS, and samples from cerebral cortices were lysed overnight in urea lysis buffer (UL) (6 M urea, 0.3 M NaCl, 1 mM EDTA, 1 mM EGTA, 25 mM TrisHCl pH 7.5, 10 mM sodium azide, 10 mM TCEP, and protease inhibitor cocktail (10 μg/mL each of Leupeptin, Pepstatin A, Bestatin, 0.8 mM Aprotinin, 1 mM AEBSF and 15 mM E-64). Lysates were cleared of precipitates at 100,000 x *g* for 1h. Protein concentration of the supernatant was determined by Bradford (BioRad). Equal volumes and concentrations of brain lysates were incubated with magnetic streptavidin beads (Pierce) on a rotator at 4^°^C overnight. Beads were washed three times with 4 M urea buffer containing 50 mM sodium phosphate and 0.1% sodium dodecyl sulfate (SDS), and three times without SDS. Biotinylated proteins were eluted by boiling the beads with LDS sample buffer supplemented with 2 mM biotin, followed by western blot analysis. The antibodies used in western blotting are listed in the Key Resource Table.

### Purifications of biotinylated proteins from mouse cerebral cortices

Mice were injected with biotin as described above. For total extract purifications, cerebral cortices were dissected from perfused brains, lysed overnight with 4 mL Neuronal Protein Extraction Reagent (N-PER, Thermo Fisher Scientific) per cerebral cortex, spun at 100,000g for 1h, and supernatants were diluted 1:1 with UL buffer before measuring protein concentrations. Next, 1 mL (750 µg/mL) of cortical lysates were treated with 20 mM iodoacetamide (Sigma-Aldrich) for 30 minutes in UL buffer, and reactions were quenched by adding DTT to 50 mM final concentration. Samples were incubated with 75 µL magnetic streptavidin beads overnight, and beads were washed as described above. After the final two washes with PBS, dry beads were flash frozen in liquid nitrogen before processing for TMT and mass spectrometry.

For membrane protein purifications, biotin-injected mice were transcardially perfused with PBS, and dissected cortices were homogenized in Dounce homogenizer with homogenization buffer containing 5 mM EDTA, 3 mM EGTA, 250 mM sucrose, 10 mM Tris-HCl pH 7.6, 1 mM phenylmethylsulfonyl fluoride (PMSF, Sigma-Aldrich), and protease inhibitor cocktail as described above. Samples were spun at 500 x *g* for 5 minutes, and supernatants were then centrifuged at 100,000 x *g* for 1 hr. P100 pellets were resuspended in WGA lysis buffer containing 150 mM NaCl, 5 mM EDTA, 3 mM EGTA, 20 mM HEPES pH 7.4, 4 mg/mL dodecyl-β-maltoside (DDM, Sigma-Aldrich), and protease inhibitor cocktail. Dissolved samples were incubated with WGA beads (Vector Laboratories) overnight. The next day, proteins bound to the WGA beads were eluted with UL buffer, and samples were processed for streptavidin enrichment as described above.

### Proteomic profiling

Dry beads were processed for TMT-MS^3^ as previously described^43,112^ by the Thermo Fisher Scientific Center for Multiplexed Proteomics (TCMP) at Harvard Medical School for the total extract experiment ➊ and the membrane protein experiment ➌ or by the Kalocsay lab for total extract experiment (Ascend instrument) ➋. Data in **Supplementary Figures S4C** were recorded on an OE480 instrument. In brief, samples were processed with the following steps:

#### On Bead Digestion and tandem mass tag (TMT) Labeling

Protein-coated beads were washed three times with PBS. On-bead digestion was performed for 3 hours at 37°C under gentle agitation with endoproteinase LysC (Wako), followed by overnight digestion with 1 µg trypsin (Promega) in 200 mM EPPS buffer (pH 8.5) containing 2% acetonitrile. The missed trypsin cleavage rate, determined by mass spectrometry (MS), remained below 10%. Acetonitrile was added to achieve a final concentration of 30% before labeling each sample with 62.5 μg of TMT 18-plex reagents (TMTPro; Thermo Fisher Scientific, Cat. #A52045). The labeling reaction proceeded for 1 hour at room temperature with intermittent vortexing. Labeling efficiency exceeded 95%, as measured by N-terminal TMTpro-conjugated peptides (dynamically searched for TMTpro [+304.207 Da] modification at the peptide N-terminus and as a static modification on lysine residues). TMT labeling reactions were quenched by adding hydroxylamine to a final concentration of 0.5% for 15 minutes, followed by acidification with formic acid. The entire volume from each sample was then pooled, and acetonitrile was evaporated to near-dryness by vacuum centrifugation (speedvac). The multiplexed sample was fractionated using the Pierce High pH Reversed-Phase Peptide Fractionation Kit (Thermo Fisher Scientific) under a modified step gradient of increasing acetonitrile (10%, 12.5%, 15%, 17.5%, 20%, 25%, 30%, 35%, 40%, 50%, 65%, 80% v/v), generating 12 fractions. Fractions were dried, desalted by StageTip, eluted with 95% acetonitrile and 1% formic acid into individual MS autosampler vials (Thermo Scientific), dried by speedvac, and finally reconstituted in 5% acetonitrile with 5% formic acid for MS analysis. Samples (Kalocsay lab) were subjected to alkaline reversed phase chromatography using the high-pH reversed phase peptide fractionation kit (Thermo Scientific #84868) and combined into 6 fractions or 12 fractions.

#### Liquid chromatography and tandem mass spectrometry

(TCMP)—Mass spectrometric data were collected on an Orbitrap Eclipse mass spectrometer coupled to a Proxeon NanoLC-1000 UHPLC (Thermo Fisher Scientific). The 100 µm capillary column was packed in-house with 35 cm of Accucore 150 resin (2.6 μm, 150Å; ThermoFisher Scientific). Data was acquired for 180 min per run. A FAIMS device was enabled during data collection and compensation voltages were set at -40V, -60V, and -80V^113^. MS1 scans were collected in the Orbitrap (resolution – 60,000; scan range – 400-1600 Th; automatic gain control (AGC) – 4×105; maximum ion injection time – automatic). MS2 scans were collected in the Orbitrap following higher-energy collision dissociation (HCD; resolution – 50,000; AGC – 1.25×105; normalized collision energy – 36; isolation window – 0.5 Th; maximum ion injection time – 100 ms.

(Kalocsay lab)—Mass spectrometric data were collected on an Orbitrap Exploris 480 or Ascend mass spectrometer coupled to a VanquishNeo UHPLC (Thermo Fisher Scientific). The 75 µm capillary column was packed in-house with 35 cm of Accucore 150 resin. Data was acquired for 60 min (OE480, 6 fractions) or 90 min (Ascend, 12 fractions) per run. A FAIMS ProDuo2 device was enabled during data collection and compensation voltages were set as described above. MS1 scans were collected in the Orbitrap (resolution – 120,000; scan range – 350-1400 Th; automatic gain control (AGC) – 4×105; maximum ion injection time – automatic). MS2 scans were collected in the Orbitrap following higher-energy collision dissociation (HCD; resolution – 50,000; AGC – 1.25×105; normalized collision energy – 33; isolation window – 0.7 Th; maximum ion injection time – 96 ms (OE480) or 400 ms (Ascend).

#### Mass Spectrometry Data Analysis

Database searching included all entries from the mouse UniProt Database (downloaded in June 2024). The database was concatenated with one composed of all protein sequences for that database in the reversed order^114^. Raw files were converted to mzXML, and monoisotopic peaks were re-assigned using Monocle^115^. Searches were performed with Comet^116^ using a 20-ppm precursor ion tolerance and fragment bin tolerance of 0.02 Da. TMTpro labels on lysine residues and peptide N-termini (+304.207 Da), as well as carbamidomethylation of cysteine residues (+57.021 Da) were set as static modifications, while oxidation of methionine residues (+15.995 Da) was set as a variable modification. Peptide-spectrum matches (PSMs) were adjusted to a 1% false discovery rate (FDR) using a linear discriminant after which proteins were assembled further to a final protein-level FDR of 1% analysis^117^. TMT reporter ion intensities were measured using a 0.003 Da window around the theoretical m/z for each reporter ion. Proteins were quantified by summing reporter ion counts across all matching PSMs. More specifically, reporter ion intensities were adjusted to correct for the isotopic impurities of the different TMTpro reagents according to manufacturer specifications. Peptides were filtered to exclude those with a summed signal-to-noise (SN) < 200 across all TMT channels and < 0.7 precursor isolation specifically. The signal-to-noise (S/N) measurements of peptides assigned to each protein were summed (for a given protein). All channels that pass QC checks with reproducibility are used for hierarchical clustering.

#### Hierarchical clustering

Proteomic results were then subjected to hierarchical clustering using JMP Pro 17 software. Candidate proteins clustered with known cilia proteins were retrieved (Tier 3) and filtered by statistical significance (p<0.05) by comparing peptide abundance between *Arl13b^+/GFP-TurboID^* samples and *Arl13b^+/+^* samples (Tier 2). Only proteins detected with more than one peptide were included in our final tier (Tier 1) as our cilia proteome.

### Expansion microscopy of tissues

Expanded mouse cerebral cortexes were prepared as described^58^. In brief, mice were perfused with PBS, followed by 4% PFA with 20% acrylamide, and 70 µm sections were cut on a VT1000S vibratome (Leica). Sections were incubated in inactivated first expansion gel solution containing 19% sodium acrylate, 10% acrylamide, and 0.1% N, N’-(1,2-dihydroxyethylene)bisacrylamide (DHEBA) in 1X PBS for 30 minutes. Next, samples were incubated in fresh, activated first expansion gel solution containing 0.075% tetramethylethylenediamine (TEMED) and ammonium persulfate (APS), and transferred to gelation chambers with 170 µm initial gel thickness. Samples were then placed in a humidified chamber, purged with nitrogen, and incubated at 37°C for 2 hr. The tissue-gel hybrids containing the region of interest were excised, and samples were incubated in denaturation buffer containing 200 mM SDS, 200 mM NaCl, and 50 mM Tris in Milli-Q ultrapure water at 37°C for 15 minutes and 75°C for 4 hr. Samples were then washed in PBS and expanded in ultrapure water overnight.

The next day, samples were re-embedded in the second gelling solution containing 10% acrylamide, 0.05% DHEBA, 0.05% TEMED, and APS in a nitrogen-filled humidified chamber at 37°C for 1.5 hr. Samples were stored in 1X PBS overnight.

On the third day, samples were incubated in the third hydrogel solution containing 9% sodium acrylate, 10% acrylamide, 0.1% N, Nʹ-methylenebis(acrylamide), 0.05% TEMED, and APS, and tissue-gel hybrids were transferred to a N_2_-filled humidified chamber at 37°C for 1.5 hr. To dissolve the DHEBA gel from the second gelation step, samples were incubated in 200 mM NaOH for 1h and washed with PBS. Samples were then incubated with primary antibodies (1:250 dilution) for 2 days. Samples were then incubated with goat anti-rabbit ATTO 647N-conjugated (Rockland #611-156-122) and goat anti-mouse IgG ATTO 550-conjugated (sigma-Aldrich #43394) secondary antibodies (1:250) for another 2 days.

On the 7^th^ day, tissue-gel hybrids were incubated in 30 µg/ml ATTO 488-conjugated NHS ester (Sigma-Aldrich #41698) for 2 hr, washed in PBS, and expanded in ultrapure water overnight.

Finally, expanded tissues were mounted on #1.5 glass-bottom dishes (MatTek) using Z-Dupe Duplicating Silicone Set (Henry Schein). Image stacks of expanded cerebral cortexes were acquired at 0.5 to 1 µm interval with 1024×1024-pixel resolution using an AX/AX R NSPARC (Nikon) with a 60x/NA 1.2 water-immersion objective. 50 to 200 z-sections were acquired to capture the entire length of a given cilium.

### Measurement of expansion factor

The linear expansion factor in expanded samples was determined as previously described^93^. Briefly, expanded samples from mouse cerebral cortices were stained with Sytox Green (Thermo Fisher Scientific), and imaged on an AX/AX R NSPARC microscope (Nikon). Images were processed with Gaussian blur, a 4-pixel Maximum filter with threshold set using the Li method. Individual nuclei were identified and segmented using MorphoLibJ^118^, measuring the cross-sectional area at the mid-plane of nuclei with circularity values greater than 0.9. In parallel, unexpanded samples from PFA-fixed cerebral cortices were stained with Hoechst. Nuclei were segmented in 3D using TrackMate^119^ with StarDist^120^ in FIJI. 3D image stacks were treated as time series to enable 2D StarDist segmentation and nuclei were reconstructed in 3D. After initial segmentation, masks were relabeled and small objects (<25,000 voxels) were removed using a custom FIJI script and MorphoLibJ. Morphological characteristics of segmented nuclei were measured using MorphoLibJ as previously described^93^. Segmented nuclei were then filtered based on sphericity (sphericity ≥0.6) using a custom Python script and visually inspected to confirm the accuracy of segmentation. Poorly segmented objects were manually removed using MorphoLibJ. The cross-sectional area of each nucleus was then measured as previously described^93^. Finally, the average mid-plane cross-sectional area from expanded nuclei was divided by that from unexpanded nuclei, and a linear expansion factor of 16.23X was applied to all pan-ExM images.

### 3D segmentation of neuronal cilia and synapses

We employed NIS-Elements (Nikon) to quantify the distance of GluN1 spots from neuronal cilia to all synapses within a given imaging volume. Using the General Analysis 3 (GA3) platform, we manually created a binary mask for cilia based on thresholding, filtered objects by size, and then segmented the final cilia via dilation, hole-filling, and surface smoothing. Neuronal synapses (using pan-stain channel) were segmented similarly, whereas GluN1 spots were segmented with a Simple Threshold function and subsequently dilated after removing granular background. We identified ciliary GluN1 spots (GluN1^cilia^) by intersecting segmented GluN1 spots with the cilia mask, and synaptic GluN1 (GluN1^syn^) by selecting GluN1 spots overlapping the segmented synapses. Child-to-Parent comparisons were then performed to measure the shortest distance from GluN1^syn^ (child) to GluN1^cilia^ (parent). As a control, the same parameters were reapplied to the same imaging volume after rotating the cilium by 90°.

3D rendering of cultured neuronal cilia (**Figs. 6K, S8C, S8F**) and brain cilia as well as synapses in pan-ExM-t images (**Fig. 7F**) was performed with Imaris 10 (Oxford Instruments) following the manufacturer’s instructions. In brief, 3D Cilia surface plots were defined by threshold of absolute fluorescent intensity and the voxel representing neuronal cilia were selected. Similarly, the segmented spots or surface of candidate proteins were using threshold methods while voxels within the surface of segmented neuronal cilia (distance between segmented proteins to cilia = 0) were selected. Surface model was used to segment all neuronal cilia and candidate proteins in *Bbs4^-/-^* neurons, while spots model was applied to candidate proteins in *Bbs4^+/-^* neurons.

### Statistical Analysis

Statistical analyses were performed using Prism 10 (GraphPad). Data from more than 2 experimental groups were analyzed by one-way ANOVA or Kruskal-Wallis test, followed by Bonferroni or Dunn post-hoc test. Data from 2 groups were analyzed using unpaired *t*-test or Mann-Whitney U test. We considered *p* < 0.05 as statistical significance. *: *p* < 0.05; **: *p* < 0.01; ***: *p* < 0.001; ****: *p* < 0.0001.

### Visualization and Gene Ontology analysis

We used Prism 10 software (GraphPad) to generate volcano plots and employed online resources, including g:Profiler (https://biit.cs.ut.ee/gprofiler/gost), CZ CELL X GENE Discover (https://cellxgene.cziscience.com), STRING (https://string-db.org), Organelle IP (https://organelles.sf.czbiohub.org), SynGO (https://www.syngoportal.org), and DISGENET (https://disgenet.com) for proteomic analysis.

## Notes

### Competing Interest Statement

The authors have declared no competing interest.

