## Supplementary material for "*In situ* Proteomics Unveils Specialized Domains for Extrasynaptic Signaling on Neuronal Cilia": Table S8

**Table S8-1**

| Alt-R® crRNA | Purpose | Sequence |
| --- | --- | --- |
| Arl13b | Making the Arl13b^+/GFP-TurboID^ mice | 5'-/AltR 1 /rArArG rGrArC rCrArA rCrCrC rArUrCrCrGrA rCrUrG rUrUrU rUrArG rArGrC rUrArU rGrCrU /AltR2/-3' |
| Arl13b V358A | Making the Arl13b^+/V358A-GFP-TurboID^ mice | 5’-/AlTR1/rUrArGrArCrUrCrGrUrCrUrGrUrArUrUrGrArCrGrUrUrUrUrArGrArGrCrUrArUrGrCrU/AlTR2/-3’ |

**Table S8-2**

| ssDNA | Purpose | Sequence |
| --- | --- | --- |
| GFP-TurboID | Making the Arl13b^+/GFP-TurboID^ mice  Note:  Homologous arms  GFP sequence  TurboID sequence  Mutated point (prevent Cas9 cleavage) | GGTTTCCATTTCCCTCCTCTGTGCAGATTTCTATGGGAAGCCGCTGCCTCCCCTGGCTGTGCGACAGAGACCTAACGGTGATGCTCAGGACACGATCTCAtccggtggctccggaccggtcgccaccATGGTGAGCAAGGGCGAGGAGCTGTTCACCGGGGTGGTGCCCATCCTGGTCGAGCTGGACGGCGACGTAAACGGCCACAAGTTCAGCGTGTCCGGCGAGGGCGAGGGCGATGCCACCTACGGCAAGCTGACCCTGAAGTTCATCTGCACCACCGGCAAGCTGCCCGTGCCCTGGCCCACCCTCGTGACCACCCTGACCTACGGCGTGCAGTGCTTCAGCCGCTACCCCGACCACATGAAGCAGCACGACTTCTTCAAGTCCGCCATGCCCGAAGGCTACGTCCAGGAGCGCACCATCTTCTTCAAGGACGACGGCAACTACAAGACCCGCGCCGAGGTGAAGTTCGAGGGCGACACCCTGGTGAACCGCATCGAGCTGAAGGGCATCGACTTCAAGGAGGACGGCAACATCCTGGGGCACAAGCTGGAGTACAACTACAACAGCCACAACGTCTATATCATGGCCGACAAGCAGAAGAACGGCATCAAGGTGAACTTCAAGATCCGCCACAACATCGAGGACGGCAGCGTGCAGCTCGCCGACCACTACCAGCAGAACACCCCCATCGGCGACGGCCCCGTGCTGCTGCCCGACAACCACTACCTGAGCACCCAGTCCGCCCTGAGCAAAGACCCCAACGAGAAGCGCGATCACATGGTCCTGCTGGAGTTCGTGACCGCCGCCGGGATCACTCTCGGCATGGACGAGCTGTACAAGtccggactcagatctcgagctcaagcttcgaattctgcagtcgaCAAAGACAATACTGTGCCTCTGAAGCTGATCGCTCTCCTGGCTAATGGCGAGTTCCATAGTGGCGAACAGCTGGGAGAAACCCTGGGCATGTCCAGGGCCGCTATCAACAAGCACATTCAGACTCTGCGCGACTGGGGCGTGGACGTGTTCACCGTGCCCGGAAAGGGCTACTCTCTGCCCGAGCCTATCCCGCTGCTGAACGCTAAACAGATTCTGGGACAGCTGGACGGCGGGAGCGTGGCAGTCCTGCCTGTGGTCGACTCCACCAATCAGTACCTGCTGGATCGAATCGGCGAGCTGAAGAGTGGGGATGCTTGCATTGCAGAATATCAGCAGGCAGGGAGAGGAAGCAGAGGGAGGAAATGGTTCTCTCCTTTTGGAGCTAACCTGTACCTGAGTATGTTTTGGCGCCTGAAGCGGGGACCAGCAGCAATCGGCCTGGGCCCGGTCATCGGAATTGTCATGGCAGAAGCGCTGCGAAAGCTGGGAGCAGACAAGGTGCGAGTCAAATGGCCCAATGACCTGTATCTGCAGGATAGAAAGCTGGCAGGCATCCTGGTGGAGCTGGCCGGAATAACAGGCGATGCTGCACAGATCGTCATTGGCGCCGGGATTAACGTGGCTATGAGGCGCGTGGAGGAAAGCGTGGTCAATCAGGGCTGGATCACACTGCAGGAAGCAGGGATTAACCTGGACAGGAATACTCTGGCCGCTACGCTGATCCGAGAGCTGCGGGCAGCCCTGGAACTGTTCGAGCAGGAAGGCCTGGCTCCATATCTGCCACGGTGGGAGAAGCTGGATAACTTCATCAATAGACCCGTGAAGCTGATCATTGGGGACAAAGAGATTTTCGGGATTAGCCGGGGGATTGATAAACAGGGAGCCCTGCTGCTGGAACAGGACGGAGTTATCAAACCCTGGATGGGCGGAGAAATCAGTCTGCGGTCTGCCGAAAAGTAATcaagtcggatgggttggtcctttttatatcagcaaggtgaactgagacctttccccaaagcagaaaagccctgaatgctgatgatgggcaagacaacca |
| Arl13b V358A | Making the Arl13b^+/V358A-GFP-TurboID^ mice | GTTCACTTTTTCAAGATATAAAGTTTCATACCTATATTCTTCTAGAAAACAGTAAGAAGAAAACCAAGAAACTACGAATGAAAAGGAGTCATCGGGCAGAACCAGTGAATACAGACGAGTCTACTCCAAAGAGTCCCACGCCTCCCCAACCTCCCCCTCCTGGTGAGTACGTCCATCCTG |

**Table S8-3**

| **Genotyping primer** | **Sequence** |
| --- | --- |
| Forward | 5’-GCG ACA GAG ACC TAA CGG TG-3’ |
| Reverse | 5’-TCG ATG TTG TGG CGG ATC TT-3’ |
| Wildtype (WT) | 5’-TGT CCA AAA GTC GGG ATG TGT-3’ |
